# Dissecting telomere maintenance mechanisms in pediatric glioblastoma

**DOI:** 10.1101/129106

**Authors:** Katharina I. Deeg, Inn Chung, Alexandra M. Poos, Delia M. Braun, Andrey Korshunov, Marcus Oswald, Nick Kepper, Sebastian Bender, David Castel, Peter Lichter, Jacques Grill, Stefan M. Pfister, Rainer König, David T. W. Jones, Karsten Rippe

## Abstract

Pediatric glioblastoma (pedGBM) represent a highly malignant primary brain tumor with recurrent mutations in the chromatin remodeler ATRX and the histone variant H3.3 that is typically associated with a fatal outcome. ATRX acts as suppressor of the alternative lengthening of telomeres (ALT) pathway, which is frequently activated in pedGBM. However, telomere features of pedGBMs have not been studied in detail, and ALT-positive model cell lines are lacking. Here, we systematically characterized a panel of pedGBM models that carry a representative set of recurrent genomic mutations for a variety of telomere features. These included the presence of ALT-associated promyelocytic leukemia nuclear bodies and C-circles, a specific type of extrachromosomal telomeric repeats, the telomere repeat content, and phosphorylation of histone H3.3 at serine 31. From an integrated analysis of seven pedGBM cell lines and 57 primary tumor samples we identified cell lines and tumors that represent the different telomere maintenance mechanisms and conclude the following: (i) A positive signal in the C-circle assay is a reliable ALT marker. (ii) ALT features occur heterogeneously and one pedGBM subgroup uses a ‘non-canonical’ ALT mechanism in the presence of wild-type ATRX. (iii) The spreading of H3.3S31 phosphorylation during mitosis is associated with loss of ATRX but not with ALT *per se*. (iv) In contrast to a previous study in glioma stem cells, we did not find a hypersensitivity of ALT cells towards the ATR inhibitor VE-821. (v) ALT-positive pedGBMs can be reliably identified from a classification scheme developed here that evaluates various combinations of cytogenetic and/or genomic data. Thus, our findings elucidate further details of the ALT pathway in pedGBMs, provide valuable models for evaluating ALT targeted therapies in a preclinical setting, and introduce an ALT classification scheme for primary tumor samples.

## Introduction

Cancer cells acquire a telomere maintenance mechanism (TMM) to avoid cellular senescence and apoptosis induced by the replicative shortening of their chromosome ends. Frequently, telomerase is reactivated to extend the telomeres. However, in 5-25 % of cases (depending on tumor entity), alternative lengthening of telomeres (ALT) pathways exist that operate via DNA repair and recombination processes [1-3]. In comparison to other tumor entities, the ALT phenotype has an unusually high prevalence in glioblastoma (GBM) [4-6]. With an estimate of 44% of ALT-positive cases, as determined by telomere FISH, pediatric GBMs (pedGBMs) are even more prone to develop ALT than adult GBMs (14%) [6]. The ALT phenotype shows a strong correlation with mutations in the DNA helicase and chromatin remodeler *ATRX* in pedGBMs (14-35%) as well as in adult GBMs (7%) [1, 5]. Interestingly, *ATRX* mutations often co-occur with mutations in *TP53* [7, 8]. Based on these recurrent mutations and on a number of mechanistic cell line studies, it is emerging that ATRX acts as tumor suppressor and inhibits the emergence of ALT as discussed in several recent reviews [9-11]. This activity appears to be related to the proper deposition of the histone variant H3.3 at heterochromatic genomic regions such as telomeres, pericentromeres, endogenous retroviral repeats, and imprinted genes. Interestingly, pedGBMs frequently harbor mutations in H3.3, namely K27M or G34R/V, which are almost exclusively found in glial tumors of childhood but it is unclear if H3.3 mutations and the ALT phenotype are linked.

Identifying the TMM that is active in a tumor is important as it can provide valuable prognostic and potentially predictive information in some cancer types [4, 12-15]. Furthermore, deregulated TMM factors represent potential targets for anticancer therapies that are mostly unique to cancer cells. While therapies that target ALT are lacking, small molecule telomerase inhibitors are currently being tested in clinical trials [16-18]. However, treatment with telomerase inhibitors may select for the emergence of an ALT-positive tumor population [19-21]. The presence of ALT is typically inferred from a number of characteristics that include very heterogeneous telomere lengths [2, 22], ultra-bright telomere foci detected by telomere FISH in tumor tissues [1], increased telomeric recombination [23], the presence of a specific type of circular mostly single-stranded C-rich extrachromosomal telomeric repeats (C-circles) [2, 24-26] and complexes of PML nuclear bodies with telomeres, termed ALT-associated promyelocytic leukemia (PML) nuclear bodies (APBs) [23, 27, 28]. More recently, two additional features have been linked to the presence of ALT: High levels of the telomeric repeat-containing non-coding RNA TERRA, and the above-mentioned mutations in the chromatin remodeler ATRX [1, 5, 29, 30]. Furthermore, the phosphorylation of serine 31 on H3.3 has been reported to spread abnormally along mitotic chromosomes specifically in ALT-positive cells [31].

Despite the high prevalence of ALT in pedGBMs, a systematic characterization and evaluation of ALT is currently lacking. It is unclear which combination of markers is suitable for reliable identification of ALT in this entity, and what would be an applicable workflow to be used in a clinical routine procedure. Furthermore, ALT has not been characterized in cell lines derived from pedGBMs and only two adult glioblastoma cell lines with an ALT phenotype have been described to date [32, 33]. Thus, studies on the ALT mechanism in pedGBMs that require *in vitro* models cannot be conducted. To address these shortcomings, we here performed a comprehensive analysis of characteristic ALT features in pedGBM tumor samples from the ICGC PedBrain cohort as well as a panel of pedGBM cell lines. These cell lines can be used to evaluate potentially ALT-specific treatment approaches, such as the application of the ataxia telangiectasia- and RAD3-related (ATR) protein inhibitor, which has previously been reported to specifically target cells that have ALT [15]. Finally, we provide a classification scheme to predict the presence of ALT in a tumor sample from a subset of measured ALT features.

## Results

### Identification of five ALT-positive pedGBM cell lines reveals a ‘non-canonical’ ALT phenotype

We analyzed seven pedGBM cell lines including the three well-established cell lines SF188, SJ-G2 and KNS42 as well as cell lines derived from freshly resected H3.3-K27M mutant pediatric high-grade gliomas (NEM157, NEM165, NEM168) and a H3.3-G34R mutant tumor (MGBM1) that were described previously [45, 46]. Previous studies have reported various mutations in these cell lines. These included a C250T mutation in the promoter of the *TERT* gene encoding the protein component of telomerase and a hypomorphic *ATRX* point mutation in the KNS42 cell line as well as severe mutations in *ATRX* in SJ-G2, MGBM1 and NEM157 [43, 47]. We additionally performed whole-genome sequencing (WGS) of the NEM165 and NEM168 cell lines and found no mutation in *ATRX* but mutations in *TP53* (Fig 1A). The panel of pedGBM cell lines covered different combinations of recurrent mutations known to occur in this entity, i.e. mutations in both *ATRX* and in the H3.3 encoding gene *H3F3A* (MGBM1, NEM157), or only in *ATRX* (SJ-G2) or only in *H3F3A* (KNS42, NEM165, NEM168). Additionally, ATRX expression was assessed by immunofluorescence. While the ATRX protein was undetectable in the severely *ATRX*-mutated cell lines, it was present in the other cells and localized in distinct nuclear foci (Fig 1A, S1 Fig). These foci largely colocalized with PML nuclear bodies as reported previously for HeLa cells and human fibroblasts [48, 49].

**Fig 1.**
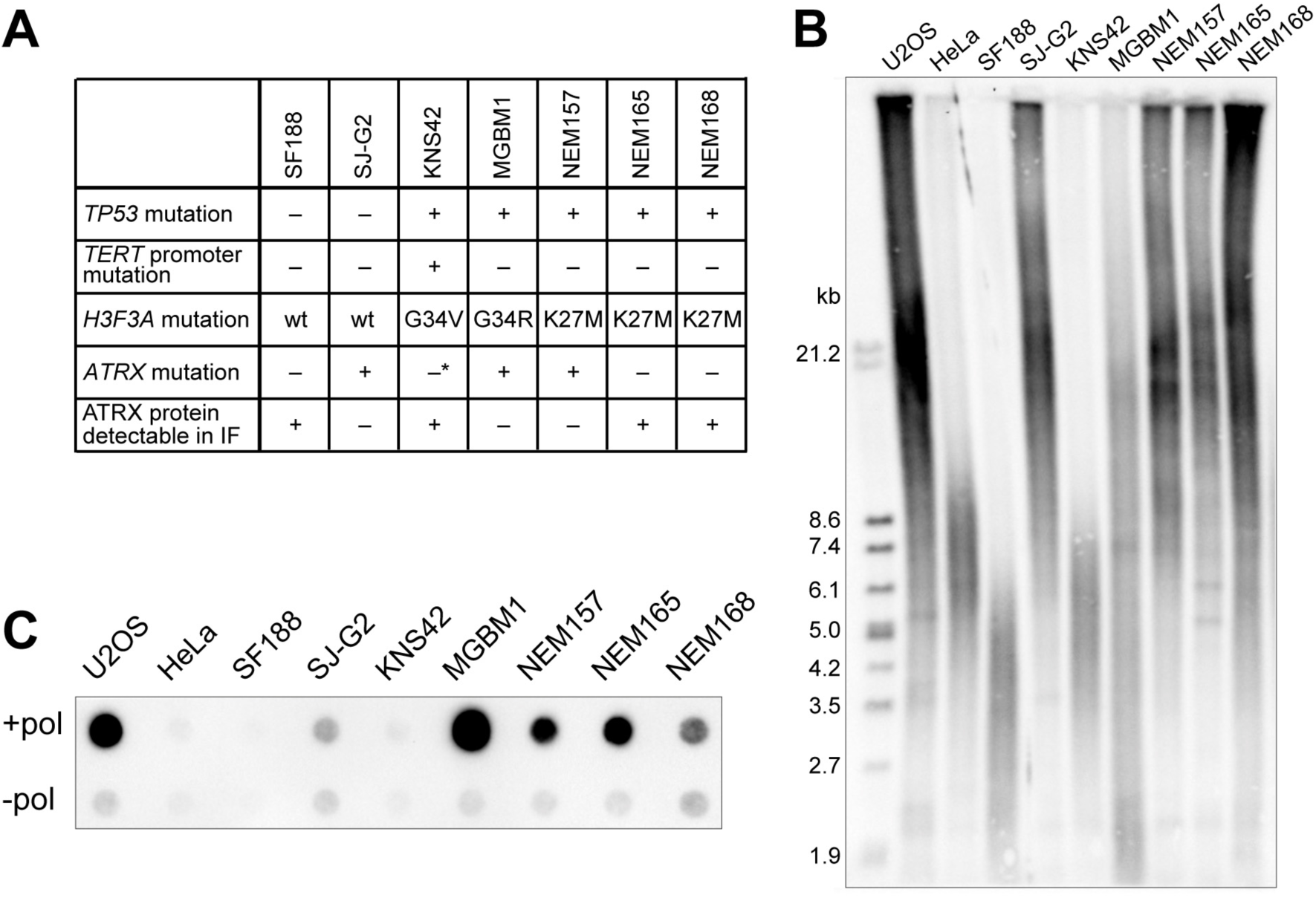
Identification of ALT-positive pedGBM cell lines. (A) Genetic mutations were determined by WGS data analysis in this study (NEM165 and NEM168) or previously in [43] (SF188, SJ-G2, KNS42, MGBM1 and NEM157). The “+” and “-” symbols illustrate the presence or absence of the indicated mutation, respectively. ATRX protein expression was detected by IF, see also S1 Fig. (B) TRF analysis visualizing telomere length distributions in pedGBM and reference cell lines (U2OS as ALT-positive, HeLa as telomerase-positive). (C) Dot-blot of C-circle assay with pedGBM and reference cell lines. For each cell line one reaction without and one with addition of polymerase (“-pol” and “+pol”) was performed.

To classify the cell lines according to their TMM, we performed terminal restriction fragment (TRF) analysis to visualize the telomere length distribution in each cell line. C-circle assays were done to assess the presence of ALT-specific extrachromosomal telomeric repeat structures. As references, the ALT-positive U2OS cell line and the telomerase-positive HeLa cell lines were included in the analyses. In the TRF blot, SF188 and KNS42 cells showed a homogenous distribution of telomere lengths characteristic of telomerase-positive cell lines, as exemplified by the reference HeLa cell line (Fig 1B). In contrast, ALT-positive tumors or cell lines typically displayed a sustained smear on the TRF blot that ranges from less than 2 kb to more than 50 kb indicating a heterogeneous distribution of telomere lengths as exemplified by the ALT-positive U2OS cell line. It was observed for SJ-G2, MGBM1, NEM157, NEM165 and NEM168 (Fig 1B).

Those five cell lines were also found to be positive in the C-circle assay, albeit with strong variations regarding the levels of these extrachromosomal telomeric repeat structures, while the SF188 and KNS42 cells were negative in the C-circle assay (Fig 1C). Based on the TRF and C-circle analysis and the TERT expression levels, SJ-G2, MGBM1, NEM157, NEM165 and NEM168 were identified as ALT-positive (Table 1). Notably, two of these, namely NEM165 and NEM168, have wild-type *ATRX* and showed normal protein expression and localization (Fig 1A, S1 Fig). This finding is particularly interesting, since the vast majority of ALT cell lines harbor *ATRX* mutations and/or show an aberrant ATRX protein expression [30]. Additionally, ATRX has been shown to act as an ALT suppressor [50, 51]. Since mutations in the ATRX binding partner DAXX have also been associated with ALT [1], we further examined *DAXX* in the NEM165 and NEM168 cell lines but did not find any mutations. Thus, we propose that the NEM165 and NEM168 cell lines employ an ALT pathway in the presence of functional ATRX/DAXX, which is in line with a recent study that suggested a high fraction of tumors with neither *TERT* activation nor *ATRX/DAXX* mutations [52]. Since both of the ALT-positive, *ATRX* wild type cell lines harbored the H3.3-K27M mutation, we further investigated whether stable ectopic expression of H3.3 mutants in SF188, SJ-G2 and HeLa LT (long telomeres) cells is sufficient to induce or enhance the ALT phenotype. The HeLa LT cell line was selected because for these cells a successful induction of ALT has been reported previously [53]. Long-term ectopic expression of H3.3 mutants was not sufficient to induce or enhance an ALT phenotype as judged from the C-circle assay (S2 Fig).

**Table 1.** TMM characteristics of pedGBM cell lines.

|  | SF188 | KNS42 | SJ-G2 | MGBM1 | NEM157 | NEM165 | NEM168 |
| --- | --- | --- | --- | --- | --- | --- | --- |
| Chromothripsis | + | + | + | – | – | + | + |
| *TP53* mutation | – | + | – | + | + | + | + |
| *TERT* promoter mutation | – | + | – | – | – | – | – |
| *H3F3A* mutation | wt | G34V | wt | G34R | K27M | K27M | K27M |
| *ATRX* mutation | – | –* | + | + | + | – | – |
| ATRX protein detectable in IF | + | + | – | – | – | + | + |
| Heterogeneous telomere lengths<sup>a</sup> | – | – | + | + | + | + | + |
| Average number of APBs per cell | 0.5 | 0.5 | 4.3 | 8.5 | 1.4 | 0.8 | 1.7 |
| C-circle levels (a.u.)<sup>b</sup> | 0.1 | 0.1 | 1.5 | 143.2 | 9.2 | 10.6 | 2.4 |
| *TERT* expression (RPKM) | 0.09 | 1.05 | 0 | 0 | 0 | 0 | 0 |
| TERRA levels (a.u.)<sup>c</sup> | 0.1 | 0.0 | 1.0 | 0.9 | 0.3 | 0.2 | 0.2 |
| Relative telomere content (qPCR)<sup>d</sup> | 0.03 | 0.06 | 0.43 | 0.21 | 0.35 | 0.33 | 1.04 |
| Aberrant H3.3S31p | – | – | + | + | + | – | – |
| Telomere maintenance mechanism | non-ALT | non-ALT (*TERT*p mut) | ALT | ALT | ALT | ALT | ALT |

ALT, alternative lengthening of telomeres
\* KNS42 has a hypomorphic mutation in *ATRX* (Q891E) [47]
<sup>a</sup> determined by TRF analysis
<sup>b</sup> determined by C-circle assay (int(+pol) - int(-pol)/ U2OS(int(+pol)-int(-pol)) (n=2)
<sup>c</sup> determined by RTq-PCR and normalized to TERRA levels in U2OS
<sup>d</sup> telomere content to single copy gene ratio (T/S)

We next analyzed a set of pedGBM samples from the ICGC PedBrain cohort by telomere FISH and C-circle assay to determine ALT activity, and compared the results with the matched sequencing information either generated in the present study or previously reported [43] (Fig 2, S3 Fig, S1 Table). The tumor samples were classified as ALT-positive based on the C-circle assay and the presence of ultra-bright telomere foci detectable by telomere FISH (Fig 2) as described in further detail below in the context of the ALT classifier scheme. Out of the 13 ALT-positive samples, 12 were found to harbor a mutation in *ATRX*, while one sample had a wild-type *ATRX* DNA sequence (S1 Table). However, this tumor (GBM33) was negative for ATRX expression when tested by immunohistochemistry, suggesting a DNA sequence independent mechanism for deregulated protein expression. Thus, the unusual ALT subgroup identified in the cell lines with functional ATRX protein was not represented in our primary tumor sample set.

**Fig 2.**
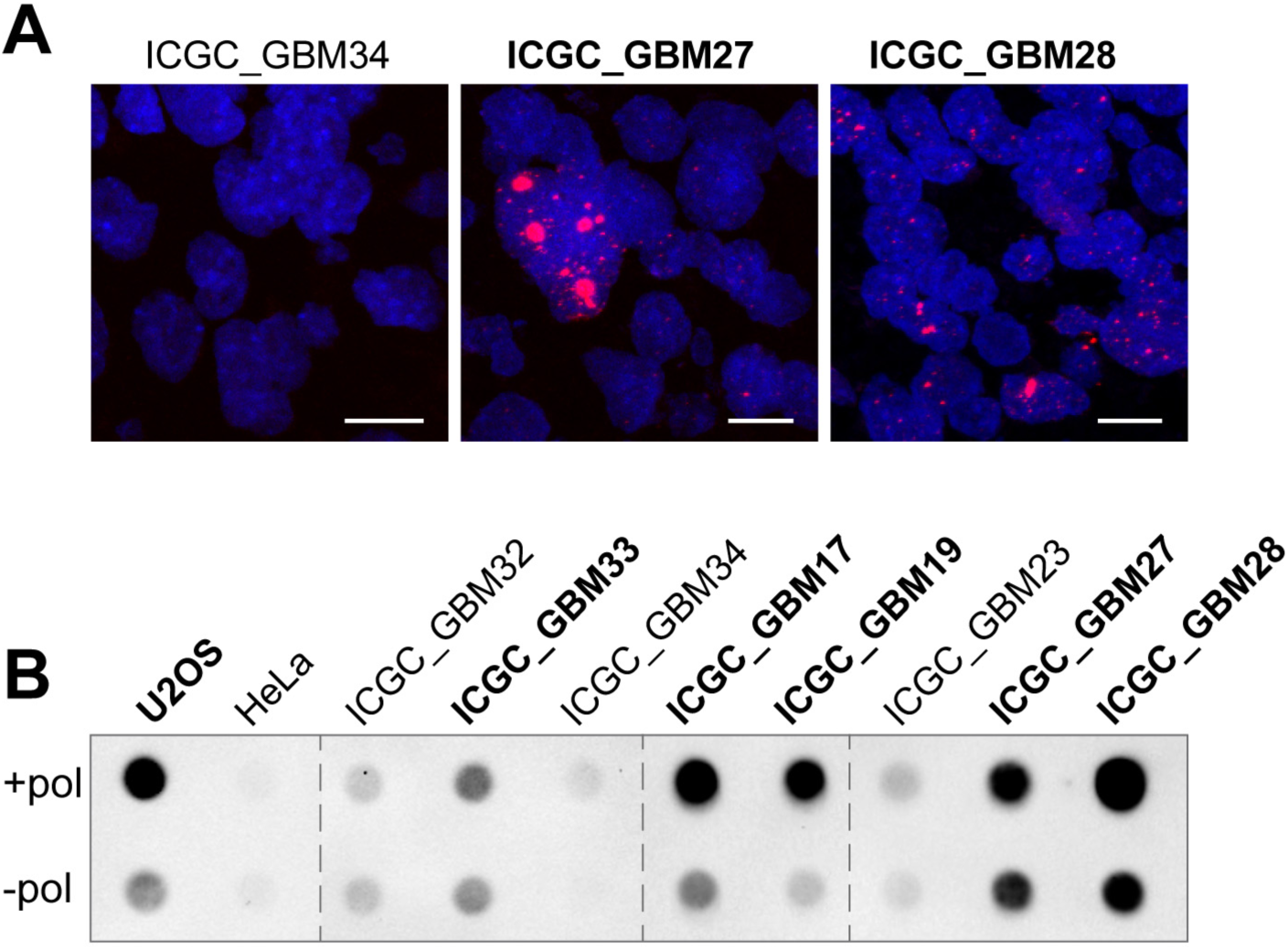
Telomere FISH and C-circle assay of primary pedGBM samples. (A) Telomere FISH was performed on pedGBM formalin-fixed and paraffin-embedded (FFPE) tissue sections using a Cy3-labeled probe targeting the C-rich telomere strand. Images were acquired by confocal laser scanning microscopy (CLSM) and maximum z-projections are shown. Exemplary samples that display ultra-bright telomere foci are indicated in bold. Scale bars are 10 μm. (B) C-circle assay of primary pedGBM samples and reference cell lines. For each sample, reactions without addition of polymerase (“-pol”) were included as controls. Samples that are classified as C-circle-positive according to the criteria described in the text are indicated in bold. Uncropped blots are shown in S3 Fig.

### ALT features are heterogeneous in ALT-positive pedGBM cell lines

A comprehensive study of ALT features in pedGBM was lacking so far. Accordingly, we characterized additional ALT features in our set of pedGBM cell lines (Fig 3) that are summarized in Table 1 together with the TMM status. The heterogeneous telomere length of ALT-positive cells evaluated in the TRF blot (Fig 1B) is frequently associated also with a higher telomere content [29]. In line with this previous observation, the telomere content of the pedGBM cell lines as determined by quantitative PCR was found to be higher in ALT-positive cells (Fig 3A). A particularly high telomere content was measured for the NEM168 cell line. Next, the presence of APBs, defined as colocalization between telomeres and PML nuclear bodies, was analyzed by telomere FISH and immunofluorescence of PML. The telomere pattern differed between the cell lines in terms of signal intensity and detectable telomere foci, again indicative of varying telomere lengths (Fig 3B). Yet, the difference in telomere lengths was much less evident than in the TRF analysis (Fig 1B). We further note that ultra-bright foci frequently detected in tissue sections of ALT-positive tumor samples are absent in cell lines. Thus, visual inspection of telomere FISH signals in cell lines is not sufficient for reliable TMM classification. For quantification of APBs, a previously established automated confocal 3D-image acquisition and analysis was employed [40]. In the non-ALT cell lines 0.5 ± 0.1 colocalizations of telomeres and PML bodies were detected per cell, which is similar to the reference telomerase-positive HeLa cell line (Fig 3C). The number of colocalizations per cell was consistently higher in the ALT-positive cell lines. These displayed a rather large variation between 0.8 APBs to 8.5 APBs per cell (Fig 3C). The latter was determined in the *ATRX* and H3.3-G34R mutated MGBM1 cell line, which thus had about twice the number of APBs as the ALT-positive U2OS reference cell line. Interestingly, the MGBM1 cell line also had a particularly high amount of C-circles (Fig 1C).

**Fig 3.**
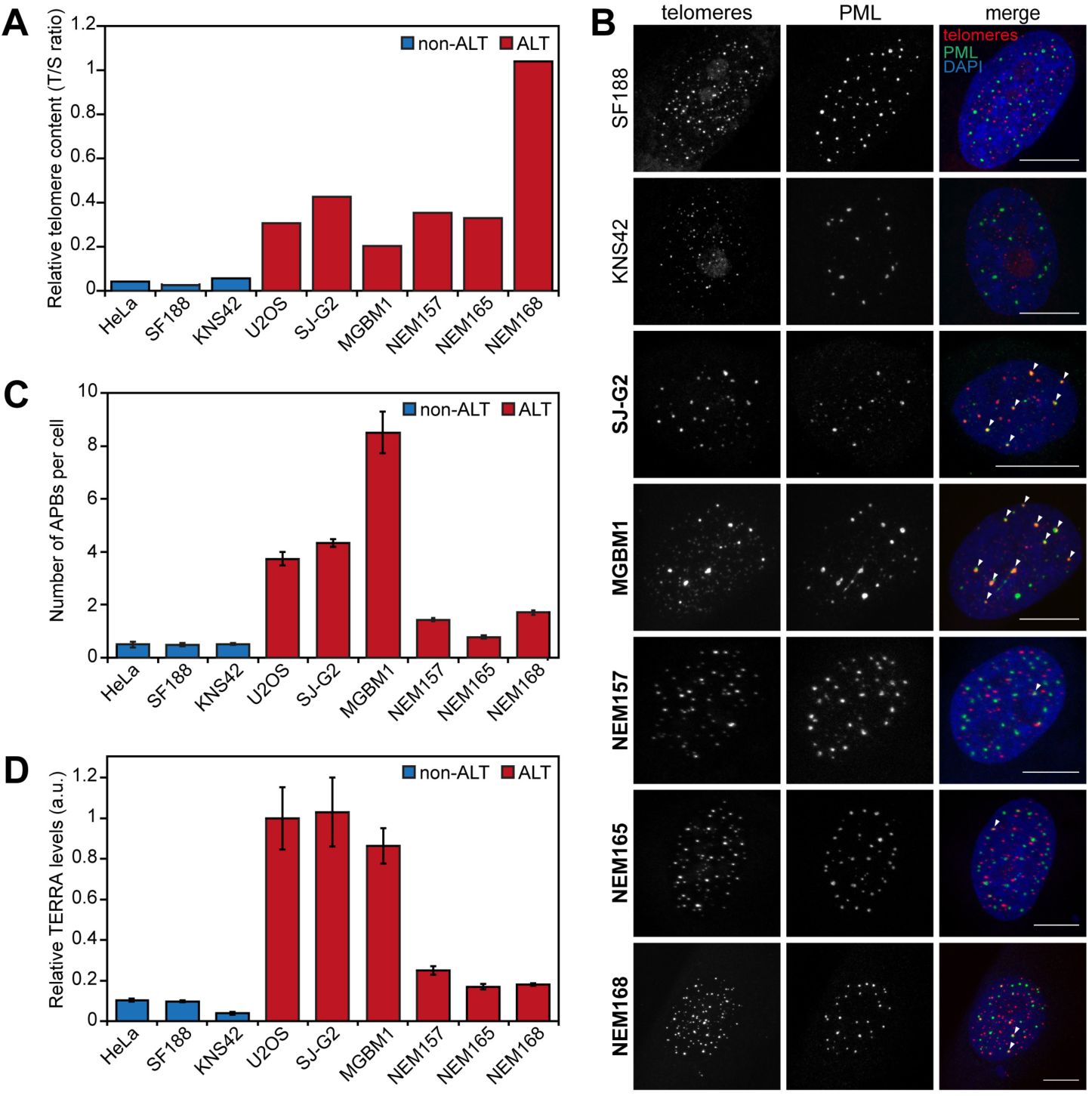
Analysis of characteristic ALT features in pedGBM cell lines. (A) Relative telomere content (T) in each sample was determined by qPCR and normalized to a single copy gene (S, *36B4*). (B) APB stainings. Representative CLSM images of pedGBM cells stained by FISH with a Cy3-labeled telomere repeat probe and by IF against PML. Colocalizations of PML and telomeres, defined as APBs, are marked by arrows in the merged images. Scale bars are 10 μm. ALT-positive cell lines are shown in bold. (C) Quantification of the number of APBs per cell based on stainings as described in panel A. Colocalizations of the telomere FISH and PML IF signals were determined by automated 3D confocal image analysis. Bars represent the average number of APBs per cell based on an analysis of at least 600 cells per cell line. (D) TERRA levels were quantified by RT-qPCR using primers that bind to the telomeric tract of TERRA that is common to all TERRA transcripts. TERRA levels were first normalized to β-actin and then to TERRA levels in U2OS. Red bars in A, C, and D represent ALT-positive cell lines as determined by TRF and C-circle analyses, blue bars represent non-ALT cell lines.

Since the telomeric non-coding RNA TERRA has been implicated in the ALT mechanism (reviewed in [11]), we next determined TERRA levels in each pedGBM cell line by quantitative reverse transcription PCR (qRT-PCR) in relation to HeLa and U2OS cells (Fig 3D). TERRA levels in the telomerase-positive cells were about 10-fold lower than in U2OS cells (Fig 3D), an ALT cell line with high TERRA levels [30]. TERRA levels in the ALT-positive SJ-G2 and MGBM1 cells were similar to U2OS. Interestingly, relative TERRA levels in the ALT-positive H3.3-K27M mutant NEM165 and NEM168 cell lines were determined to be 5-fold less than in U2OS cells, while still being 2-fold higher than in HeLa cells. This indicates that in general TERRA is a good marker for ALT. However, ALT determination solely based on TERRA levels could lead to an inaccurate classification. TERRA levels have been reported to correlate with hypomethylation of the subtelomeres, where the TERRA promoter is located [29, 54]. Accordingly, methylation at CpG sites located within one megabase (Mb) of the chromosome end was analyzed based on Illumina 450K methylation array data [39]. These regions were not found to be differentially methylated in the two H3.3-G34R/V mutant cell lines studied here (S4 Fig).

### Aberrant H3.3 serine 31 phosphorylation is linked to ATRX loss rather than ALT *per se*

A recent study reported that ALT-positive cells specifically display high levels of histone H3.3 serine 31 phosphorylation (H3.3S31p) on the entire chromosome, due to an elevated activity of CHK1 serine/threonine kinase [31]. We therefore analyzed the distribution of H3.3S31p on metaphase chromosomes of the pedGBM cell lines. For telomerase-positive pedGBM cells, H3.3S31p was confined to a region close to the centromeres, in line with previous reports [31, 55]. In contrast, the phosphorylation was spread over the entire chromosomes in *ATRX*-mutated ALT-positive cells (Fig 4). Notably, this was not the case for *ATRX* wild type ALT-positive NEM165 and NEM168 cells. In the latter cell lines H3.3S31p was mainly restricted to the centromeres and pericentromeres, resembling the H3.3S31p staining of telomerase-positive cells. A similar observation has been reported previously for the ALT-positive, *ATRX* wild type SKLU1 cell line [31]. Hence, the aberrant H3.3S31p correlates with the absence of ATRX loss rather than ALT activity. The wild-type *H3F3A* containing SJ-G2 cell line displayed the same H3.3S31 phosphorylation pattern mitotic chromosomes as the cell lines MGBM1 and NEM157, which harbor the G34R- and the K27M-mutations, respectively. Thus, the presence of *H3F3A* mutations did not show any obvious correlation with the aberrant H3.3S31p mark.

### ALT-positive pedGBM cells are not hypersensitive to ATR inhibition

An intact shelterin complex protects telomeres from being erroneously recognized by the DNA damage response machinery, including ATR protein kinase signaling. In ALT-positive cells, ATR inhibition leads to a reduction of ALT markers [15, 53]. Inhibition of ATR in combination with other treatments has also been reported to have an effect on viability of cancer cells in general [56, 57]. In addition, a hypersensitivity of ALT-positive tumors has been proposed by Flynn et al. [15]. Accordingly, we tested ATR inhibitor sensitivity for the panel of 7 pedGBM cell lines introduced here. In line with our previous study on non-glioma cell lines [42], we found no correlation of ALT status and response to the VE-821 ATR inhibitor (Fig 5A). Instead, the sensitivity varied between cell lines irrespective of the telomere maintenance mechanism. For example, the ALT cell line SJ-G2 was highly sensitive to the ATR inhibitor, whereas another ALT-positive cell line, MGBM1, was resistant to the treatment. In addition, we quantified induced cell death by FACS analysis of annexin V and propidium iodide stained ALT and non-ALT cells treated with 3 μM VE-821 for 6 days (Fig 5B, S5A Fig). To better compare the results of the cell viability assays with FACS results, we tested the influence of cell density on VE-821 sensitivity and observed that a higher starting cell density results in a lower sensitivity (S5B Fig). This is consistent with our previous observation of a strong influence of the initial cell number in this assay [42]. In general, we did not observe a selective killing of ALT cells by ATR inhibition in these experiments.

**Fig 4.**
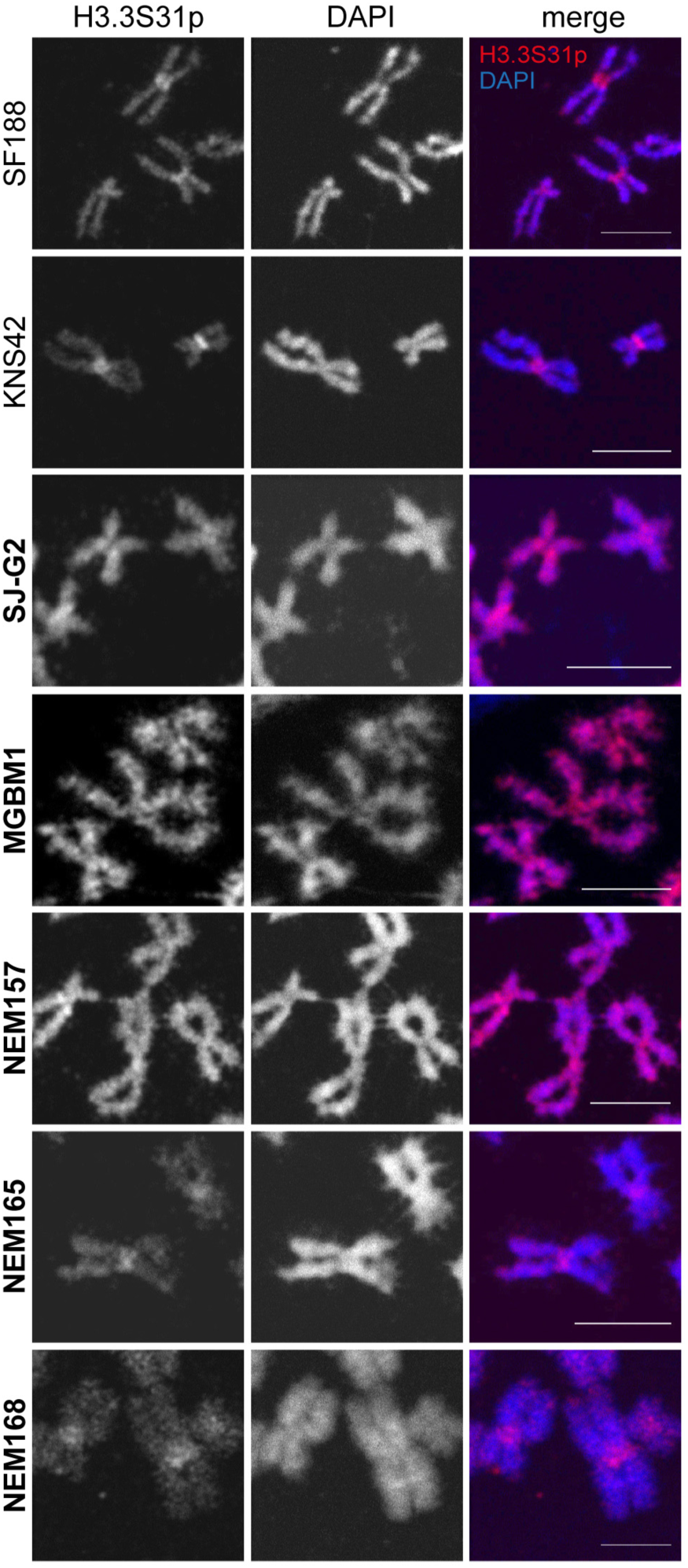
Localization of H3.3S31p on the chromosome arms in pedGBM cell lines. Immunofluorescence stainings were performed on native metaphase spreads using an antibody against H3.3S31p. Images were acquired with identical microscopy settings. Scale bars are 5 μm. ALT-positive cell lines are shown in bold.

**Fig 5.**
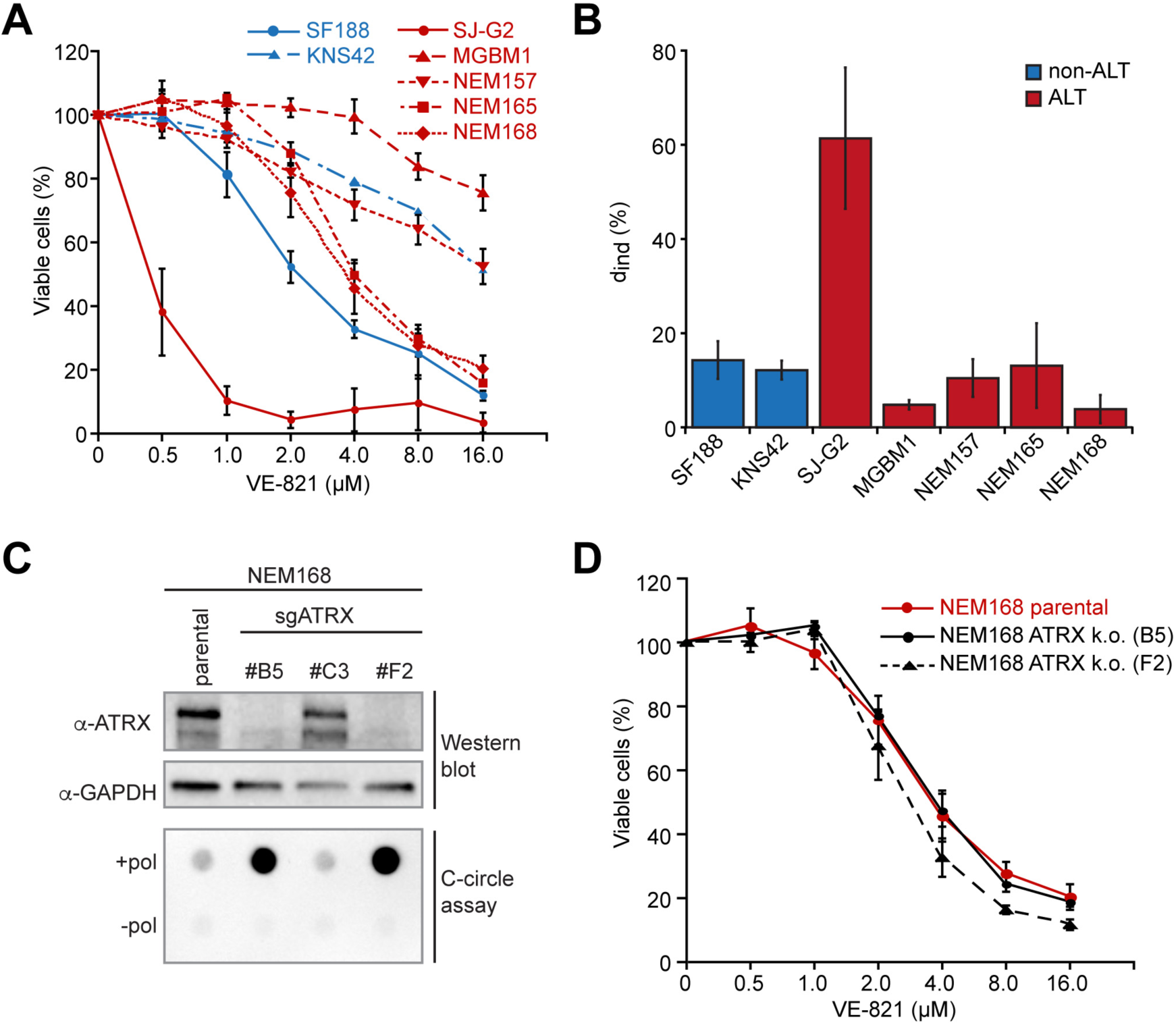
Cell viability of ALT-positive and -negative pedGBM cell lines upon treatment with the ATR inhibitor VE-821. (A) Analysis of cell viability of pedGBM cell lines in response to the ATR inhibitor VE-821. After 6 days of treatment with the indicated concentrations of VE-821 cell viability was analyzed using Celltiter Glo. Error bars represent the standard error of three experiments. (B) Percentage of induced cell death determined from FACS analysis as in S5A Fig. Induced cell death was calculated as described in the Materials and Methods section. (C) *Top,* ATRX western blot to detect CRISPR/Cas9 mediated loss of ATRX in the NEM168 clones #B5 and #F2. GAPDH was used as loading control. *Bottom,* C-circle assay of the same cell clones. Enrichment of the “+pol” over the “-pol” sample is strongly increased in the cells that carry the *ATRX* knockout. (D) Cell viability analysis of the parental *ATRX*-proficient NEM168 and the *ATRX* k.o. NEM168 cells upon treatment with different VE-821 concentrations as in panel *a.*

In order to validate this finding in an isogenic setting, we generated *ATRX* knockout cells of one of the pedGBM cell lines, NEM168, using CRISPR/Cas9. ATRX has been shown to repress the ALT phenotype upon re-introduction into ATRX-deficient, ALT-positive cells [42, 50]. Accordingly, we detected a more pronounced ALT phenotype in ALT-positive NEM168 cells after *ATRX* knockout (Fig 5C) when evaluating C-circle levels. We next tested whether this enhanced ALT phenotype affects VE-821 sensitivity, which would be expected if ALT rendered cancer cells hypersensitive towards ATR inhibitors as proposed [15]. However, the loss of ATRX and concomitant enhancement of the ALT phenotype did not result in an increased VE-821 sensitivity (Fig 5D). In summary, characteristics of the cell lines other than the TMM determine the cellular response to ATR inhibition, which confirms our previous observations in non-glioma cells [42].

### A selected set of features defines the TMM status in a pedGBM tumor cohort

We next analyzed pedGBM tumors from the ICGC cohort in more detail by integrating their previously published (epi)genetic features [43] in relation to their telomere maintenance mechanism. Based on the C-circle assay results introduced above, we classified 13 samples as ALT-positive (Fig 2B, S3 Fig, S1 Table). A positive C-circle signal is a reliable indicator for the presence of ALT [26]. However, the lack of signal in the C-circle reaction has to be interpreted with caution. The mostly single-stranded C-circles can rapidly degrade during repeated freeze-thaw cycles and may thus have been lost during storage of the tumor DNA. For a non-overlapping fraction of samples from the same cohort, telomere FISH was performed to detect ultra-bright telomere foci (Fig 2A, S1 Table). A high agreement between telomere FISH and C-circle assay was detected in our study. This finding corroborates previous observations of ultra-bright telomeric foci in tissue sections being a good ALT marker [1]. Based on these observations, we defined those samples with a positive C-circle result as ‘ALT’ and the ones that tested negative for both C-circles and ultra-bright foci as ‘non-ALT’. In addition, the C250T and C228T mutations in the *TERT* gene promoter were used as a reference feature for the telomerase-positive non-ALT phenotype. The C250T/C228T nucleotide exchanges create novel binding sites for ETS transcription factors and result in *TERT* expression [58, 59]. Of the three tumor samples that carry the C228T mutation, two were subjected to the C-circle assay and revealed no signal (S1 Table). This finding is in line with previous literature and the hypothesis that the occurrence of ALT and *TERT* promoter mutations are mutually exclusive [60].

For the pedGBMs samples (epi)genetic maps were determined previously [43]. These data were evaluated here in terms of the TMM status for our reference data set as defined above. Based on previous published literature we selected the following features as informative for a TMM classification (S1 Table): (i) Loss of function of the chromatin remodeler ATRX as inferred from sequencing analysis and/or the lack of protein expression seen in immunohistochemistry (IHC) strongly correlated with ALT. (ii) Among the 7 tumor samples that harbored mutations in the H3.3 encoding *H3F3A* gene and were analyzed with respect to their ALT status, 5 were found to be ALT-positive. (iii) In line with previous reports that associated *TP53* mutations with ALT, 12 of the 13 C-circle-positive tumors also harbored *TP53* mutations [61-63]. (iv) Chromothripsis, the mutational ‘shattering’ of large parts of a chromosome has been functionally connected to dysfunctional telomeres and ALT activity [64, 65]. In our set of reference samples we found a higher number of chromothripsis-positive samples that were non-ALT compared to ALT-positive ones. Thus, there is no association between ALT and chromothripsis. (v) *TERT* expression as detected by RNA-seq. In line with previous reports the number of *TERT* transcripts was low (S6A and S6B Figs), but their detection still correlated with the classification as ‘non-ALT’, further supporting the notion that in these samples telomerase is activated [66, 67]. (vi) The DNA methylation status of a region upstream of the *TERT* transcription start site was used as an additional surrogate marker for *TERT* expression [68]. The samples evaluated as ALT-positive showed consistently lower *TERT* promoter methylation compared to the ‘non-ALT’-samples. Thus, the latter group or part of it maintains their telomeres via a promoter DNA methylation-linked upregulation of telomerase (S1 Table, S6C and S6D Figs). (vii) As described above for the pedGBM cell lines, telomere content was usually higher in ALT-positive cells compared to ALT-negative ones (Fig 3A). Using qPCR we measured the relative telomere content in a subset of pedGBM tumor samples where tumor and matched control blood samples were available. As expected, the ‘ALT’ tumor reference samples displayed a somewhat higher telomere content ratio than the ‘non-ALT’ group (S6E and S6F Figs).

The analysis of differentially expressed genes in ALT and non-ALT tumors from RNA-seq data revealed 141 significant genes (adjusted p-value ≤ 0.05, 65 genes with adjusted p-value ≤ 0.01). However, no obvious TMM gene signature was identified that could be used for a classification (S7 Fig).

In summary, we defined a reference data set for TMM classification by evaluating samples with a positive C-circle signal as ‘ALT’ and those without C-circle signal in combination with the absence of ultra-bright foci in telomere FISH as well as samples with C250T/C228T *TERT* promoter mutations as ‘non-ALT’. Moreover, the additional features described above were selected to provide further information on the TMM status by integrating them into a classification scheme.

### The ALT status can be evaluated with a decision tree-based classifier

Next, we systematically evaluated the relation of the above-described features with ALT occurrence. Furthermore, we identified suitable combinations to predict if ALT is active in a given pedGBM sample. This analysis was conducted by developing a decision tree-based ALT classifier for pedGBM tumors. At http://www.cancertelsys.org/paint/index.html a web-based tool termed *PAINT* (Predicting Alt IN Tumors) is available. *PAINT* makes predictions from the available, frequently incomplete data set of a sample and allows the integration of results from very different techniques, i.e. DNA/RNA-sequencing, methylation array, IHC, qPCR, and FISH. For constructing such a classifier, the values of all features introduced above were translated into binary values and then used to construct decision trees as described in Materials and Methods (Fig 6A). The accuracy of the resulting decision tree is given as a performance value *P* where *P* = 1 represents 100% correct prediction with the samples from the reference data set. In addition, a *p*-value (*p*) was calculated for each tree based on the confusion matrix to test if it is better than a random one. Using each feature alone, ultra-bright telomeric foci (*P* = 0.95), ATRX protein expression (*P* = 0.90), and *ATRX* mutation (*P* = 0.89) were the best predictors for ALT in pedGBM (Fig 6A, S2 Table). On the other hand, information on chromothripsis alone is not able to predict the class reliably (*P =* 0.63, *p >* 0.05). The decision tree based only on telomere content is also not associated with a significant *p*-value (S2 Table). For the latter case, the low number of reference samples should be taken into account (n = 15), and the addition of further samples to the training set might improve predictions based on this feature in the future. The ATRX status has a strong predictive power in pedGBM and addition of further features such as *TERT* expression and telomere content improves the performance of the resulting decision tree only marginally from *P* = 0.90 to *P* = 0.91 (*p* = 3.12 x 10^−5^) (Fig 6B). On the other hand, merging features that on their own were only relatively weak predictors, e.g. *TP53* mutation (*P* = 0.69) and *TERT* expression (*P* = 0.77), or were not significant such as telomere content (*P* = 0.73, *p* = 0.14), into one decision tree can clearly improve the performance of the prediction (*P* = 0.82, *p* = 4.29 x 10^−4^) (Fig 6C, S2 Table).

**Fig 6.**
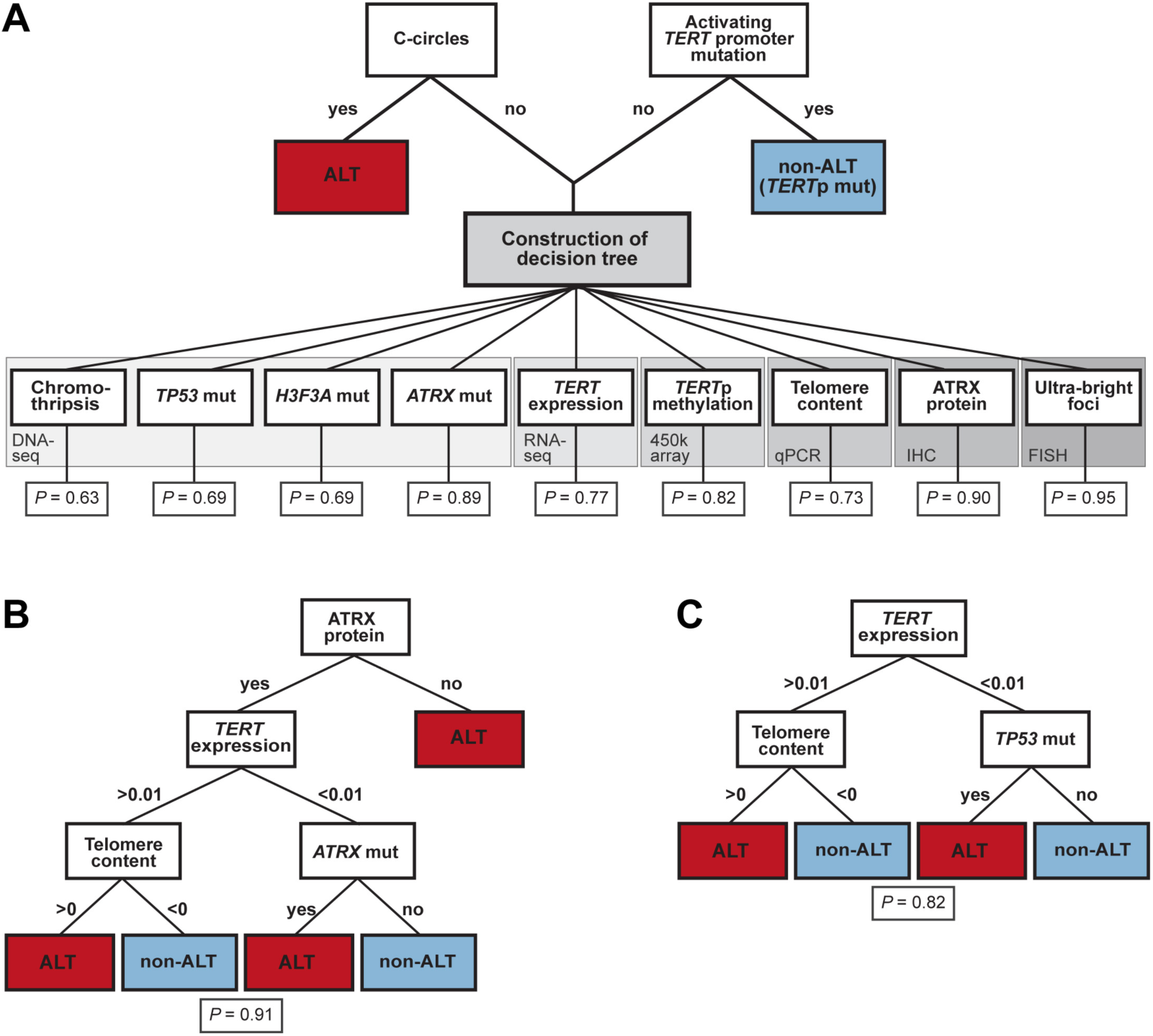
ALT classification scheme. (A) Scheme for classification of primary pedGBM samples according to their TMM status. Accumulation of C-circles is evaluated as an ALT-defining characteristic. Similarly, the presence of an activating *TERT* promoter mutation determines that a tumor is employing telomerase as the activated TMM and thus is classified as ‘non-ALT’. If none of these two criteria are true, the other parameters listed here are taken into account to classify a sample as ‘ALT’ or ‘non-ALT’. Note that the information is obtained from different types of assays that require different sources of material, such as DNA for C-circle assay, detection of mutations, methylation assays, and PCR, RNA for estimation of expression, and tissue sections for IHC and FISH procedures. The performance value *P* indicates the predictive power of the respective feature by itself as determined by the classifier. For the respective *p*-values please refer to S2 Table. (B) Decision tree using ATRX protein expression, *TERT* mRNA expression, telomere content, and *ATRX* mutation. Note that for construction of this decision tree also *TP53* mutation was part of the input feature set for training purposes, but was not used in the resulting consensus tree depicted here (*P* = 0.91, adjusted *p* = 3.12 x 10^−5^, see also S2 Table). (C) Decision tree to classify a pedGBM sample as ‘ALT’ or ‘non-ALT’ based on *TERT* expression, *TP53* mutation and telomere content. The tree with this combination of parameters predicts the presence of ALT with a performance of *P* = 0.82 (adjusted *p* = 4.29 x 10^−4^) clearly improving the prediction compared to the use of either of these features alone.

### Discussion

Patients suffering from glioblastoma have a very poor prognosis with treatment options being often limited and ineffective [4, 69]. Thus, it is important to gain a better understanding of the tumor biology underlying GBMs. Assessment of telomere maintenance mechanisms represents a novel promising approach towards this goal. The majority of (adult) GBMs employ telomerase to maintain their telomeres. However, the ALT mechanism has an unusually high prevalence in pediatric GBM, with up to 44% of ALT-positive cases, as assessed by ultra-bright foci in telomere FISH [4, 5, 70]. Yet, other ALT markers have not been studied in detail and ALT-positive pedGBM cell lines as suitable model systems were lacking. Here, five ALT-positive pedGBM cell lines were identified. These cell lines provide valuable models to gain a better understanding of the ALT pathway in pedGBMs and to test novel ALT-targeted therapies in a preclinical setting. For this purpose it is also of advantage that the identified ALT-positive cell lines have different genetic backgrounds (Table 1) representing distinct epigenetic and biological subgroups of pedGBMs [39]. The detailed analysis of characteristic ALT markers revealed that the extent to which these features are present varied considerably among the ALT-positive cell lines (Table 1). The NEM165 and NEM168 cell lines displayed all the hallmarks of ALT without carrying an *ATRX* mutation, which points to a ‘non-canonical’ ALT mechanism. It is noted that the diffuse intrinsic pontine glioma (DIPG) type of pedGBM represented by the NEM cell lines carries *ATRX* mutations less frequently than non-brainstem pGBMs [71, 72]. Interestingly, we found that removing *ATRX* from the NEM168 cell line resulted in a pronounced accumulation of C-circles indicative of increased ALT activity (Fig 5C). Thus, *ATRX* mutations might be a secondary event that prolongs and/or enhances an attenuated ALT mechanism in place initially.

In summary, we conclude that some variation of the ALT phenotype exists in pedGBM with respect to the degree to which typical ALT features are present. Another main observation from our characterization of the pedGBM cell lines refers to the aberrant pattern of H3.3 serine 31 phosphorylation, which was previously linked to the presence of ALT [31]. We show here that this modification is not a direct ALT marker but in the cell lines tested here rather is associated with the loss of ATRX. The mechanism by which this leads to an extended mitotic H3.3S31p signal is unknown. One model would be that loss of ATRX results in an increase of stalled replication forks and accumulation of single-stranded DNA. This in turn could trigger an RPA- and ATR-mediated activation of CHK1, which was implied in establishing the H3.3S31p mark [31, 73].

In contrast to a previous report, ALT activity is not suited as a biomarker to predict sensitivity to ATR inhibition, which confirmed our experiments with non-glioma cell lines [15, 42]. While drug sensitivity varied among the cell lines, no association with the active TMM was observed. Thus, an efficient drug that specifically targets ALT-positive tumors is still lacking. Nevertheless, the active tumor-specific TMM is an attractive therapeutical target that needs to be further exploited in the future. In this context, a reliable TMM classification of primary tumor samples is crucial to determine whether a patient could benefit from telomerase inhibiting agents or ALT targeting drugs. In addition, the stratification of GBM patients according to the active TMM provides valuable prognostic information as ALT is associated with longer survival [4, 61, 74]. However, as discussed above typical ALT features display a significant degree of variations in pedGBM cell lines. Thus, it is essential to base the ALT identification on a suitable set of ALT markers.

Consistent with reports for other tumor entities we find the enrichment of C-circles in pedGBM to be a very good marker for ALT, with a high agreement between this assay and the detection of ultra-bright telomere foci visualized by FISH [61]. This is of particular interest as telomere FISH currently is the major technique to detect ALT. However, it requires tissue section material, expertise in FISH and a way to quantitatively measure the telomere-specific fluorescence for a proper interpretation as discussed previously [75]. In contrast, for executing a C-circle assay as little as 30 ng of DNA is sufficient, thus making it a reasonable method for clinical applications. However, a draw-back of this assay is the instability of the mostly single-stranded C-circles since their detection can be impaired by improper storage and frequent freeze-thaw cycles and lead to false-negative results [26]. Hence, combination with other ALT markers becomes essential. Accordingly, we introduce the *PAINT* decision tree-based classifier as a web tool that integrates results from up to 11 different features to predict the presence of ALT in a given pedGBM sample (Fig 6). Importantly *PAINT* integrates sequencing-based information with results from other assays. In addition, it allows taking parameters into account, which on their own are much less conclusive with regards to the active TMM (Fig 6C). For example, the mutational status of *TP53* by itself does not provide a convincing marker for ALT. However, the TMM prediction is largely improved when combined with information about the telomere content and the RNA-seq derived *TERT* expression, which by itself is also difficult to interpret due to generally low *TERT* mRNA levels [66, 67]. It is noted that the prediction of the ALT status from sequencing data via *PAINT* can be used for a retrospective analysis of cohorts that have been sequenced previously to link the TMM with disease progression. This will largely improve patient stratification with respect to their activated TMM since much larger sample numbers are available if telomere FISH or C-circle data are not required.

In the current implementation the decision tree is based on a binary feature discrimination with continuous data as from DNA methylation arrays or RNA-seq analyses being converted into binary values. In the future, the performance of current classifiers can be improved when integrating support vector machines. Extending the reference sample set of pedGBM samples with defined ALT status and its implementation as training data in the classification scheme will also increase the prediction accuracy.

The activation of a TMM is an important step in oncogenesis and an integrated analysis of (epi)genomic features in relation to the telomere length is relevant for the analysis of all cancer types [52]. Accordingly, it will be informative to implement an expanded *PAINT* classifier approach that combines sequencing- and imaging-based TMM analysis with molecular assays into a pan-cancer tool. With a growing number of samples and more information to train the classifier we will also be able to extend the scheme and distinguish more groups such as telomerase-positive, ALT with loss of ATRX, ALT in the presence of functional ATRX, and cases that do not display any signs of both ALT and telomerase activation. The latter groups are particularly interesting as a recent study found 22% of cancers with no sign for telomerase activation or *ATRX*/*DAXX* aberrations [52]. Accordingly, we envision that the approach described in our present study will support the TMM analysis and subsequent patient stratification not only in pedGBM but also for other tumor entities that have a high prevalence of ALT such as neuroblastoma or soft tissue sarcoma [2].

## Materials and Methods

### Cell culture

U2OS, HeLa cells and HeLa LT cells were cultured in DMEM and RPMI1640 medium (Gibco), respectively, each supplemented with 10% FCS, 2 mM L-glutamine and 100 μg/ml penicillin/ streptomycin. SF188, SJ-G2, KNS42 and MGBM1 were cultured in high-glucose DMEM supplemented as described above. NEM157, NEM165 and NEM168 were cultured in Amniomax C-100 Basal Medium with 10% Amniomax C-100 supplement (both Gibco). All cell lines were cultured at 37°C in 5% CO_2._

### Generation of NEM168 ***ATRX* knockout cell lines**

Two candidate guide RNA sequences directed against *ATRX* were cloned into pX459v2.0 as described in [34]. The construct was transfected into NEM168 using TransIT-LT1 transfection reagent (Mirus), transfected cells were selected using 1.5 μg/ml puromycin and single clones were assayed for indels in the *ATRX* gene using the Surveyor assay (IDT). Deletion of *ATRX* in positive clones was validated in a western blot with anti-ATRX antibody (HPA001906, Sigma).

### Terminal restriction fragment (TRF) analysis

Genomic DNA was purified using the Gentra Puregene Cell Kit (Qiagen) and DNA integrity was assessed on a 1% agarose gel. Genomic DNA (5 μg) was digested overnight with the restriction enzymes *Hinf* I and *Rsa* I (12.5 U each). The digested DNA was resolved on a 0.6% agarose gel (gold, Biozym) in 0.5X TAE buffer using the CHEF-DRII pulsed-field gel electrophoresis system (Biorad) with the following settings: 4 V/cm, initial switch time 1 s, final switch time 6 s, and 13 h duration. As size references, a DIG-labeled molecular weight marker from the TeloTAGGG telomere length assay kit (Roche) was loaded alongside the digested DNA. Southern blotting and chemiluminescent detection was performed using the TeloTAGGG kit according to the manufacturer’s instructions. The resulting chemiluminescent signals were measured with a Chemidoc MP imaging system (Biorad).

### C-circle assay

The C-circle assay was essentially performed as described previously [26]. Briefly, genomic DNA was isolated from cell lines using the QIAamp DNA mini kit (Qiagen). 30 ng DNA (in 10 μl) was combined with 10 μl 2X Φ29 buffer supplemented with 7.5 U Φ29 DNA polymerase (New England Biolabs), 0.2 mg/ml BSA, 0.1% (v/v) Tween-20, 1 mM each of dATP, dGTP and dTTP and incubated at 30°C for 8 h, followed by an incubation at 65 °C for 20 min. Reactions without addition of the Φ29 polymerase were included as control (“–pol”). After addition of 40 μl 2x SSC, the amplified DNA was dot-blotted with a 96-well dot blotter onto a 2x-SSC-soaked nylon membrane. The membrane was baked for 20 min at 120°C and hybridized and developed using the TeloTAGGG kit according to manufacturer’s instructions. The result of the C-circle assay was evaluated as ALT-positive if the signal intensity of the reaction with polymerase (“+pol”) was at least 1.4-fold higher than the intensity of the C-circle signal of the reaction without polymerase (“-pol”) and additionally, at least fourfold higher than the background.

### TERRA qRT-PCR

Total RNA was extracted with the RNeasy kit (Qiagen) and subsequently digested with DNase I (Promega) for 30 min to ensure depletion of contaminating genomic DNA. Total RNA (1 μg) was reverse transcribed with gene-specific primers (Telo RT 5′-CCC TAA CCC TAA CCC TAA CCC TAA CCC TAA-3′, β-actin RT 5′-AGT CCG CCT AGA AGC ATT TG-3′, see ref. [35]) using Superscript III at 55 °C for 1 h, followed by RNase H treatment. Reactions without reverse transcriptase were performed as controls. qRT-PCR analysis was performed using SYBR green master mix (Roche). Reactions were set up in triplicates with 500 nM telomere specific primers (forward 5′-CGG TTT GTT TGG GTT TGG GTT TGG GTT TGG GTT TGG GGT-3′, reverse (5′-GGC TTG CCT TAC CCT TAC CCT TAC CCT TAC CCT TAC CCT-3′) as described [36]. β-actin specific primers were used as described previously [35]. The amplification program was as follows: 95°C for 10 min followed by 36 cycles at 95°C, 58°C and 72°C each for 10 s. TERRA levels were normalized against β-actin signals and a standard curve was used to obtain relative quantities of TERRA.

### Telomere qPCR

Telomere-repeat quantitative PCR was conducted essentially as described previously [36, 37]. In short, 5 ng DNA, 1x Lightcycler 480 SYBR Green master mix and 500 nM each of forward and reverse primer were added per 10 μl reaction (for telomere repeats telo-fwd 5’-CGG TTT GTT TGG GTT TGG GTT TGG GTT TGG GTT TGG GTT-3’ and telo-rev 5’-GGC TTG CCT TAC CCT TAC CCT TAC CCT TAC CCT TAC CCT-3’ and for the single copy gene *RPLP0/36B4* 36B4-fwd 5’-CAG CAA GTG GGA AGG TGT AAT CC-3’ and 36B4-rev 5’-CCC ATT CTA TCA TCA ACG GGT ACA A-3’). Cycling conditions were 10 min at 95 °C, followed by 40 cycles of 95 °C for 15 s and 60 °C for 1 min. A standard curve was used to determine relative quantities of telomere repeats (T) to those of the single copy gene (S). The log2 ratio of telomere content was determined by dividing the T/S ratio of the tumor sample by the T/S ratio of the control sample. The calculated log2 ratio represents the increase or decrease in telomere content in tumor versus control samples.

### Immunofluorescence and FISH

Immunofluorescence and telomere FISH on cell lines were performed as described previously [38]. The following antibodies were used: mouse anti-PML (1:100, Santa Cruz, sc-966) and rabbit anti-ATRX (1:500, Bethyl Labs, A301-045). Immunohistochemistry for ATRX (Sigma, HPA001906; dilution 1:750) on FFPE samples was performed as previously described [39]. Interphase telomere FISH was performed using a PNA probe kit (Dako).

### H3.3S31p staining on metaphase spreads

Cells were arrested in mitosis by adding Karyomax colcemid solution (Gibco) at a final concentration of 0.1 μg/ml to the culture medium and incubating for 2 h. Mitotic cells were harvested by shake-off and resuspended in a small volume of medium. Pre-warmed hypotonic solution (0.5% sodium citrate) was added dropwise to the cell suspension to obtain a final concentration of 2 x 10^4^ cells/ml. After incubation at 37 °C for 10 min, 500 μl of the cell suspension was cytocentrifuged on a microscopic glass slide (1200 rpm, 5 min). Slides were air-dried shortly and transferred to KCM buffer (120 mM KCl, 20 mM NaCl, 10 mM Tris-HCl pH7.2, 0.5 mM EDTA, 0.1% (v/v) Triton X-100) for 15 min at RT. Next, slides were incubated in anti-H3.3S31p rabbit antibody (Active motif, 1:100 in 10% goat serum in KCM buffer) for 1 h at room temperature in a humid chamber. Slides were washed two times for 5 min in KCM buffer and incubated with anti-rabbit Alexa 488 secondary antibody (Invitrogen, 1:300 in 10% goat serum in KCM buffer). Slides were again washed two times for 5 min in KCM buffer, fixed for 10 min in 4% (v/v) paraformaldehyde in KCM buffer, washed for 5 min in H2O, air-dried and mounted in prolong gold antifade reagent with DAPI (Thermo Fisher Scientific).

### Confocal image acquisition and analysis

Fluorescence microscopy images were acquired with a Leica TCS SP5 confocal laser scanning microscope (oil immersion objective lens, 63x, 1.4 NA) and are displayed as maximum intensity projections. For the quantification of APBs, z-stacks of cells stained for PML and telomeres were acquired by automated confocal image acquisition, as described previously [40, 41]. Colocalizations of PML and telomeres, representing APBs, were quantified using a 3D-model-based segmentation approach as described previously [40, 41].

### ATR inhibitor sensitivity assays

For cell viability assays, 1,500 cells were seeded in triplicate in 96-well plates and incubated overnight as described previously [42]. The following day cells were either treated with DMSO (control) or with increasing concentrations of the ATR inhibitor VE-821 (Selleckchem) dissolved in DMSO. Cells were incubated for 6 days without medium change and cell viability was analyzed using Celltiter Glo (Promega) and a TECAN Infinite M200 plate reader according to the manufacturers’ instructions. For FACS analysis, cells were seeded in T25 flasks. Each cell line was either treated with 3 μM VE-821 or with the same volume of DMSO for the control samples. After incubation for 6 days without medium change, cells (including dead cells) were collected by trypsin and total cell numbers were determined using the LUNA cell counter (Biozym). Cells were resuspended in FACS binding buffer (10 mM HEPES, 2.5 mM CaCl_2_, 140 mM NaCl) at a final concentration of 2x10^6^ cells/ml, stained with FITC annexin V (Biolegend) and propidium iodide (Invitrogen) according to the manufacturers’ instructions, and analyzed by flow cytometry on a FACS Canto II (BD Biosciences). Cells that were negative for both PI and Annexin V were identified as viable cells. Annexin V positive events were characterized as apoptotic cells, whereas PI positive events were labeled as necrotic. All fractions of viable, apoptotic and necrotic cells were quantified using the WEASEL software. The percentage of induced cell death *d*_ind_ was calculated as

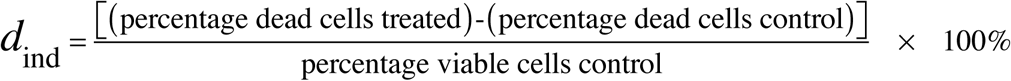

### Whole-genome sequencing (WGS) and data analysis

Whole-genome sequencing and data analysis were performed as part of the ICGC PedBrain Tumor Project, as previously described [43]. Additional new samples (ICGC_GBM84, ICGC_GBM95, ICGC_GBM96, ICGC_GBM98, ICGC_GBM100, and cell lines NEM168, NEM165) were processed in the same way.

### Accession codes

Sequencing and methylation data are available at the European Genome-phenome Archive (*http://www.ebi.ac.uk/ega/*), hosted by the European Bioinformatics Institute under the accession number EGAS00001001139 (see also [43]).

### Construction of decision tree-based classifier

To distinguish between different TMMs in pedGBM tumor samples, a classification scheme was developed. A sample with a positive C-circle signal was classified as ALT-positive, while a sample with an activating *TERT* promoter mutation was classified as ALT-negative (and *bona fide* telomerase-positive). For samples for which both features were negative or not available, a binary decision tree was constructed based on intelligent enumerating. TMM-specific features were extracted from imaging data (presence of ultra-bright telomere foci, ATRX loss of expression from immunohistochemistry staining) and combined with parameters derived from whole-genome sequencing (chromothripsis, loss of function mutations in *ATRX* and *TP53*, K27M and G34R/V mutations in *H3F3A*), RNA-sequencing (*TERT* expression), telomere qPCR (telomere content) and from methylation data from the Illumina 450K array (*TERT* promoter methylation). In total, feature information was available for seven cell lines (Table 1) and 57 pediatric glioblastoma patients (S1 Table). WGS information was mainly extracted from the data in ref. [43]. For DNA methylation as well as the telomere content, only data from primary tumor samples were used. The decision tree-based classifier was constructed from 35 samples with known TMM status. The training set consisted of the 7 cell lines (5 ALT and 2 non-ALT), 13 ALT-positive tumor samples and 15 non-ALT tumor samples, of which two were telomerase-positive as inferred from detection of a *TERT* promoter mutation. The C-circle signal determined the class ‘ALT’ (C-circle-positive) or ‘non-ALT’ (C-circle-negative and no ultra-bright FISH foci) and was not used for constructing the decision tree. The feature values from Table 1 and S1 Table were translated into binary values with 1 representing the presence of a feature. For continuous values (*TERT* promoter methylation, *TERT* expression in RPKM and telomere content), optimal thresholds were determined for ALT and non-ALT samples from the training data set. The thresholds were 0.01 RPKM for *TERT* expression (S6B Fig), a methylated fraction of 0.22 for *TERT* promoter methylation (S6D Fig), and a telomere content log 2 ratio of 0 relative to the healthy control sample (S6F Fig). Possible binary decision trees were compared to identify the tree that had the minimal number of misclassified samples from the training set and the minimal number of questions. A leave-one-out cross-validation approach was used to determine the performance of the tree, which is measured by the percentage of correct predictions in the training data and indicated by the performance value *P*. Finally, the optimal tree was derived using the complete set of samples (S2 Table). In cases where the number of samples that simultaneously had multiple features of interest was small, it was tested if the construction of the decision tree could be improved by including additional samples where a missing feature was arbitrarily selected as positive or negative. Since the number of samples with overlapping information on the features was different depending on the analyzed set of features, variations in the performance values can occur even when the same set of features was used. In cases where the performance of a tree did not improve with more features for the same number of samples,the tree with the best subset was used. In addition, a *p*-value was calculated for every tree based on the confusion matrix using Fisher’s exact test corrected according to Benjamini and Hochberg [44]. The *p*-value indicates if the calculated confusion matrix is better than a random one. We note that non-significant *p*-values in most cases are due to a small sample size for training the respective tree.

The decision trees constructed with this approach were applied to predict the class (‘ALT’ or ‘non-ALT’) of the 29 samples with unknown TMM status. A total of 12 samples were predicted as ALT and 17 as non-ALT (S1 Table). It is noted that the input is calculated for any (incomplete) combination of known features from the corresponding decision tree.

Finally, the method was implemented as a web-based program termed *PAINT* for Predicting ALT IN Tumors. It is available at *http://www.cancertelsys.org/paint/index.html* to predict if a sample is ALT-positive or -negative. The user can select a combination of available information for a given pedGBM sample and the predicted ALT state is returned. To assess the prediction quality, *PAINT* also displays the performance *P*, the *p*-value as well as the sample size used to construct the corresponding tree (see S2 Table). In cases of non-significant performances (adjusted *p*-value > 0.05), *PAINT* replaces the feature set with the subset that has a significant *p*-value (adjusted *p*-value ≤ 0.05) and the best performance.

## Acknowledgments

We thank Caroline Bauer, Stefan Wörz, Karl Rohr, the DKFZ Genomics and Proteomics Core Facility, and the DKFZ Flow Cytometry Core Facility for help and support.

## SUPPLEMENTARY INFORMATION

### Dissecting telomere maintenance mechanisms in pediatric glioblastoma

Katharina I. Deeg, Inn Chung, Alexandra M. Poos, Delia M. Braun, Andrey Korshunov, Marcus Oswald, Nick Kepper, Sebastian Bender, David Castel, Peter Lichter, Jacques Grill, Stefan M. Pfister, Rainer König, David T.W. Jones, and Karsten Rippe

### Supplementary Tables

**S1 Table.**
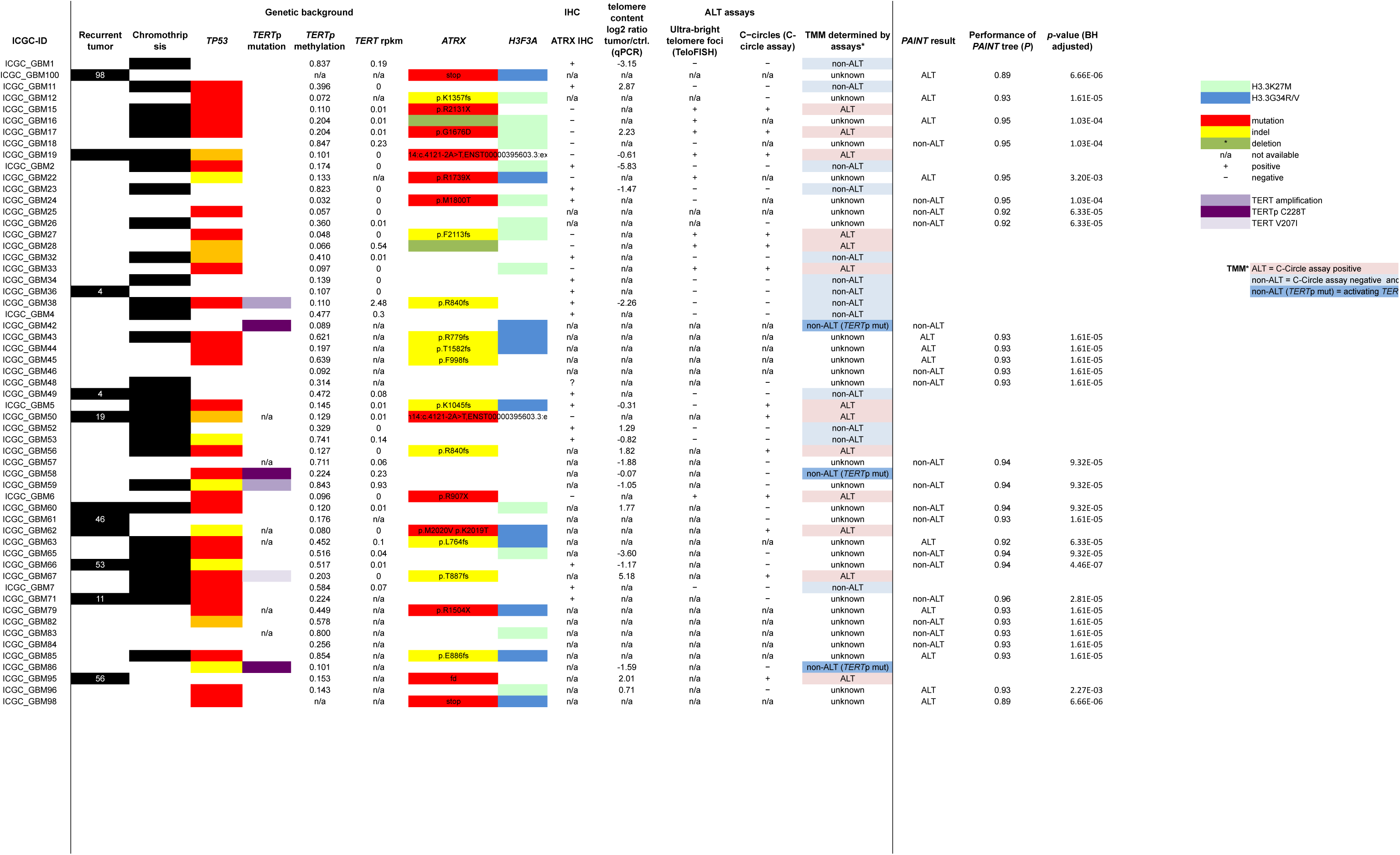
(provided as separate Excel file). Overview of genetic background, ATRX IHC, telomere content, and ALT features in pedGBM samples. Overview of pedGBM samples analyzed in this study. Sequencing data were obtained from the work of the ICGC PedBrain consortium [1]. When available, matched tissue sections were subjected to ATRX IHC and/or telomere FISH and isolated DNA was used for the C-circle assay and telomere-specific qPCR. For further analysis we defined two classes: ‘ALT’ includes samples that displayed positive C-circle signals. The ‘non-ALT’ class contains all samples, for which both of the ALT-specific assays telomere FISH and C-circle assay gave negative results, as well as tumors with an activating *TERT* promoter mutation. These classes were used to train the *PAINT* classifier as described in the text. The predicted class for those samples that did not fall into the training set is reported (column “*PAINT* result”). In addition, the performance *P* and the adjusted *p*-value of the calculated tree used for the prediction are listed in the last two columns.

**S2 Table (provided as separate Excel file). Combinations of parameters used in the *PAINT* classifier and their predictive power.**

For all possible combinations from 1 up to 9 features (“Input feature set initially tested”) the best tree was calculated by minimizing the number of misclassified samples from the training set and the number of questions. The performance of each tree was determined using a leave-one-out cross-validation, which gives rise to the performance value *P*. In addition. a *p*-value was calculated based on the confusion matrix to test if the confusion matrix is better than a random one (last column, *p*-values were adjusted using the method of Benjamini and Hochberg (BH)) [3]. Since not all samples were subjected to all assays, the information about the features was not complete for all samples. Therefore, the trees for each feature combination were calculated with different sample sizes, which introduced performance variations. Especially trees with a low sample size (n < 12) frequently have a non-significant *p*-value. If a tree with only a feature subset yielded a better performance than the tree constructed with all selected features using the same set of samples, the tree from the subset was taken as indicated in the column “Feature set selected to construct decision tree”. In cases where the sample size was small due to unavailable feature information, a decision tree was constructed by including additional samples where a missing feature was arbitrarily selected as positive or negative. If the performance of this optimized tree was better than the reference tree (only samples with complete information), it was selected (marked as ‘yes’ in column “improved *P* with n/a feature information”). Accordingly, the same best tree for different combinations of input features can exist. Together, this indicates which combinations of features can predict ALT best, but also if testing an additional parameter would improve the predictions.

## Supplementary Methods

### Viral transduction

Histone H3.3-overexpressing SF188 cells and H3.3 variant-containing constructs pLVX-Puro_H3.3-wt_HA, pLVX-Puro_H3.3-K27M_HA, pLVX-Puro_H3.3-G34R_HA have been described in ref. [2]. Lentiviral particles were produced in HEK293T cells the supernatant containing the lentiviral particles was used for transduction of SJ-G2 and HeLa LT cells. Transduced cells were selected by adding puromycin to the medium in a final concentration of 3 μg/ml.

### RNA-Seq

Total RNA was purified and genomic DNA was removed as described above. Ribosomal RNA was removed with the Ribozero gold kit (Illumina) following the manufacturer’s instructions. RNA libraries were prepared using the NEB next ultra directional RNA library preparation kit according to the manufacturer’s protocol for use with ribosome depleted RNA. Library quality was assessed on a Bioanalyzer using an Agilent High Sensitivity Chip. Two independent RNA libraries were prepared from each cell line and sequenced on a HiSeq 2000, single-read, 50 bp). The reads from RNA-seq were mapped to the UCSC hg19 assembly allowing for no mismatches using TopHat. Expression levels of *TERT* were determined in reads per kilobase per million mapped reads (RPKM).

### Analysis of differentially expressed genes

After prediction of the presence of ALT using the decision tree-based classifier all samples from ref. [1], for which RNA-Seq data was available, were used to derive differential expressed genes between ALT and non-ALT samples. Differential expression was calculated with DESeq2 [5] based on the raw read counts of the samples. For calculating the significance, DESeq2 employs the Wald test and adjusts for multiple testing using the method by Benjamini and Hochberg [3].

### DNA methylation profiling

For genome-wide assessment of DNA methylation, pedGBM cell lines and tumor samples were arrayed using the Illumina human methylation 450 bead chip according to the manufacturer’s instructions at the DKFZ. For addressing subtelomeric methylation of cell lines, the IDs of all CpG sites that were located within 1 Mb of the chromosome ends were determined using R and the length of each chromosome as specified in the UCSC hg19 assembly. The median methylation value of the selected CpG positions was determined for each cell line.

### Supplementary Figures

**S1 Fig.**
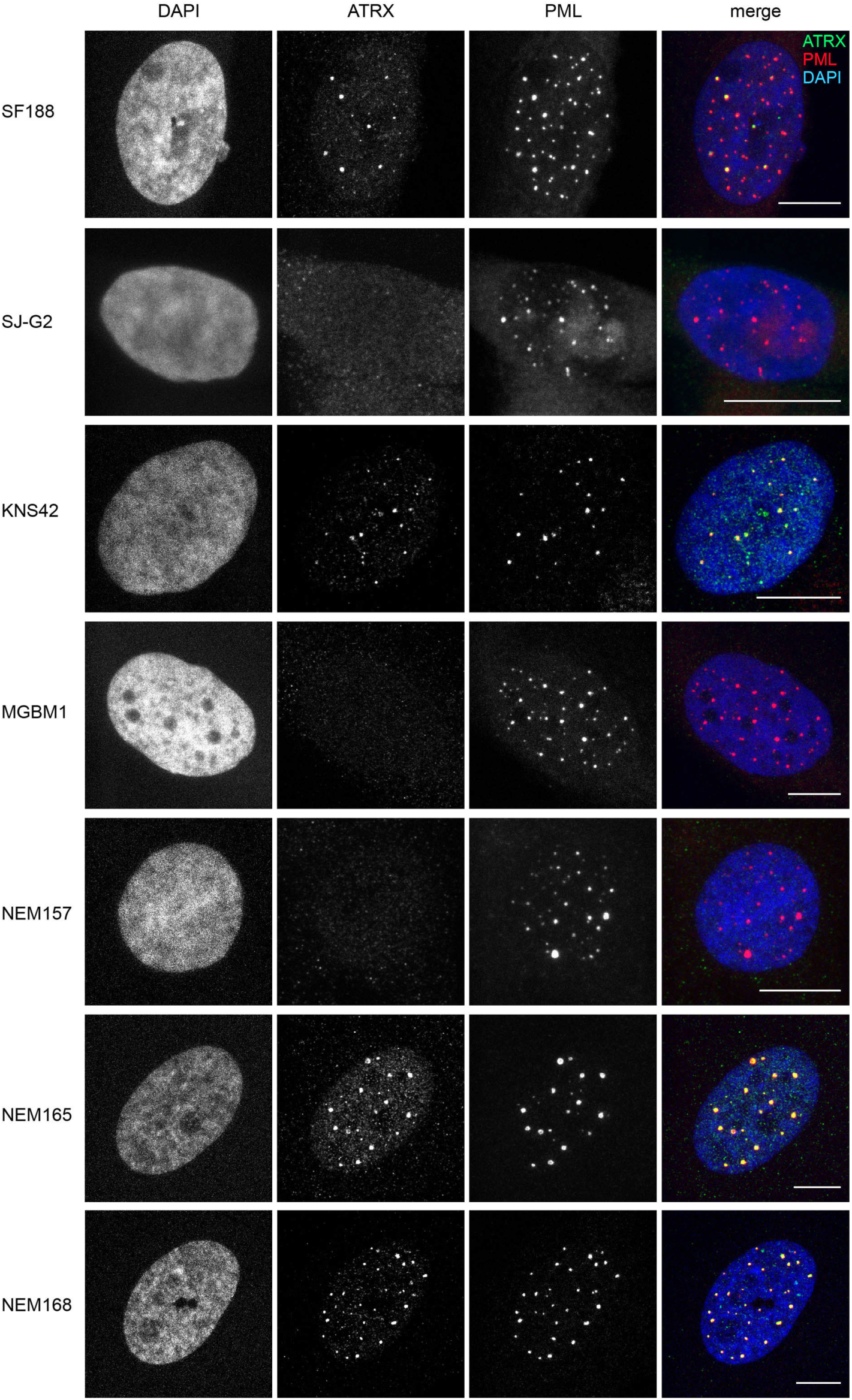
Immunofluorescence stainings of ATRX and PML in pedGBM cell lines. Maximum intensity projections of confocal microscopy image stacks acquired from ATRX and PML immunofluorescence stainings in the indicated cell lines are shown. Scale bars are 10 μm.

**S2 Fig.**
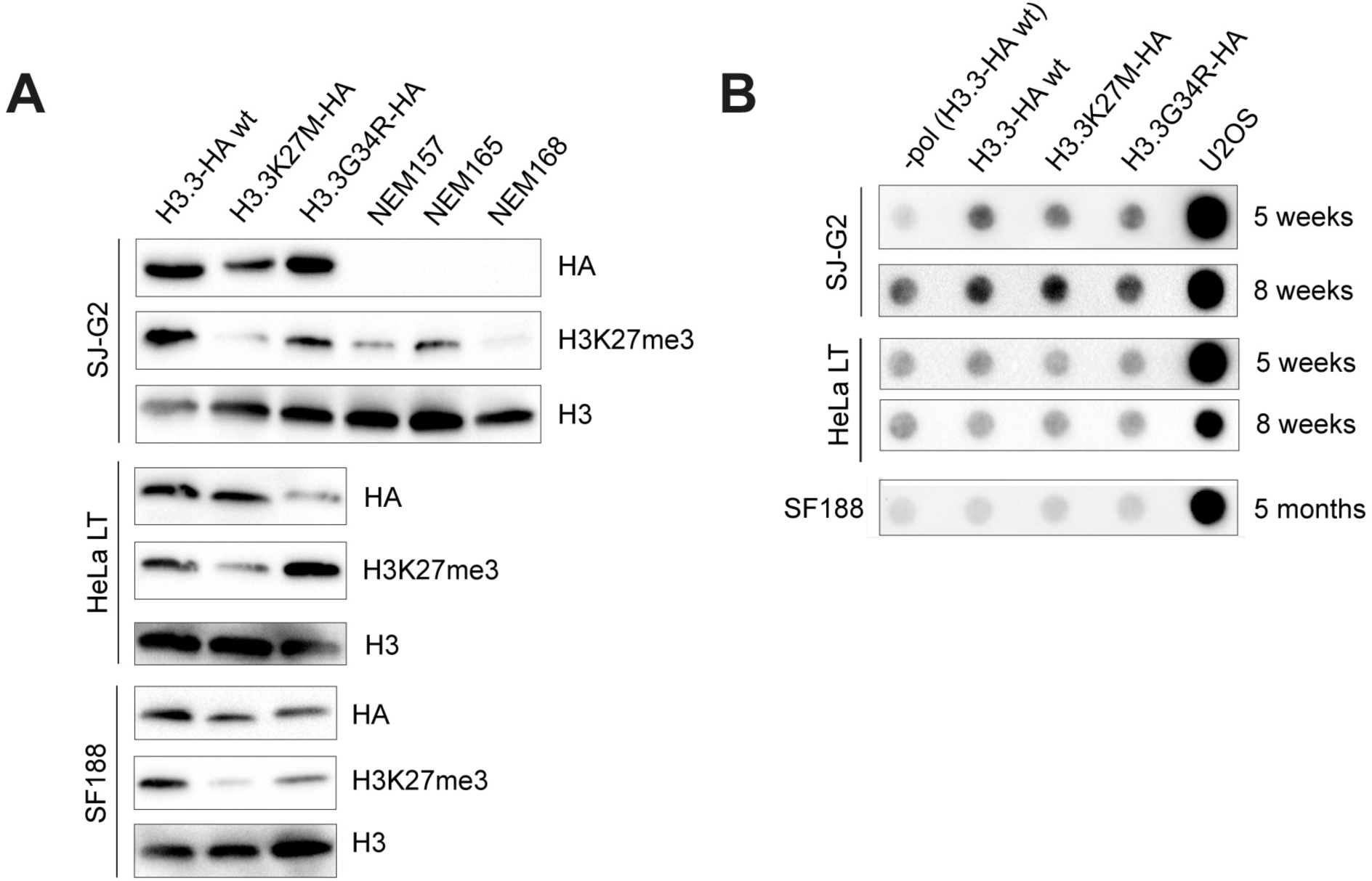
Western blots and C-circle assays of SJ-G2, HeLa LT and SF188 cells that ectopically express HA-tagged H3.3 variants. (A) Western blot analysis of SJ-G2, HeLa LT and SF188 cells stably expressing HA-tagged H3.3 variants (wildtype, wt; K27M; G34R). Histones were isolated 8 weeks (SF188, HeLa LT) or 5 months (SF188) after lentiviral transduction and expression of HA-tagged constructs was analyzed by Western blot using an anti-HA antibody. H3K27me3 levels were visualized by an anti-H3K27me3 antibody. Histone H3.3 or histone H3 expression was used as a loading control. (B) C-circle levels upon ectopic expression of H3.3 mutant proteins in SJ-G2, HeLa LT and SF188 cells. ALT-positive U2OS cells were included as reference. Genomic DNA was isolated at the indicated time points after lentiviral transduction and C-circle assay was performed with 30 ng DNA. A reaction without addition of polymerase (“-pol”) was included as control.

**S3 Fig.**
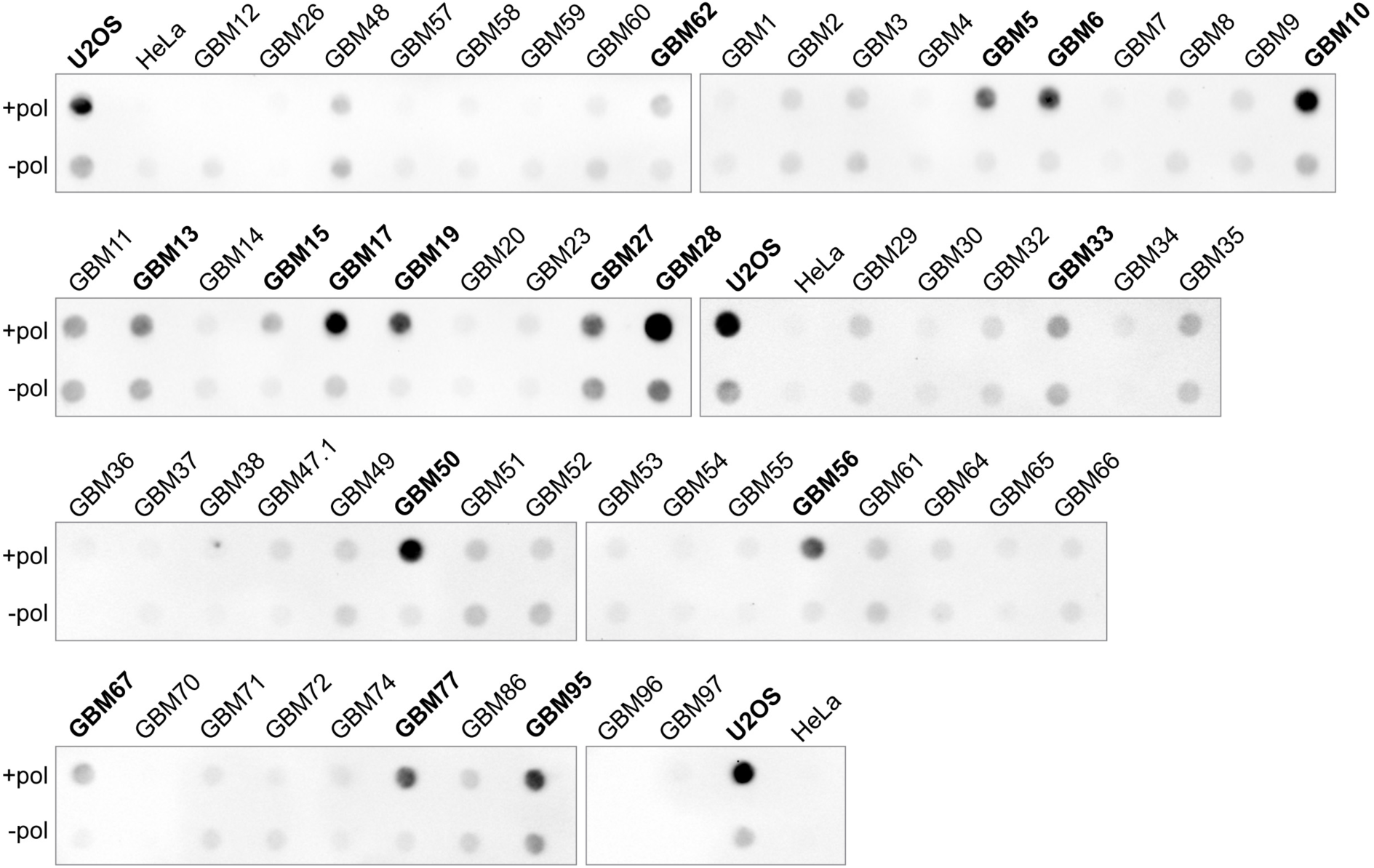
C-circle assay of pedGBM tumor samples. Dot-blot of C-circle analysis with telomerase-positive HeLa cells and ALT-positive U2OS cells as references. For each sample, reactions without addition of polymerase (“-pol”) were included as controls. Samples that are classified as C-circle-positive according to the criteria described in the text are indicated in bold. Parts of the C-circle blots presented here are also shown in Fig 2. Note that not all of the samples depicted here appear in S1 Table. Some samples were excluded from the final analysis due to too low tumor content or identification as a tumor entity different from GBM. Exposure time for all blots shown was 240 s.

**S4 Fig.**
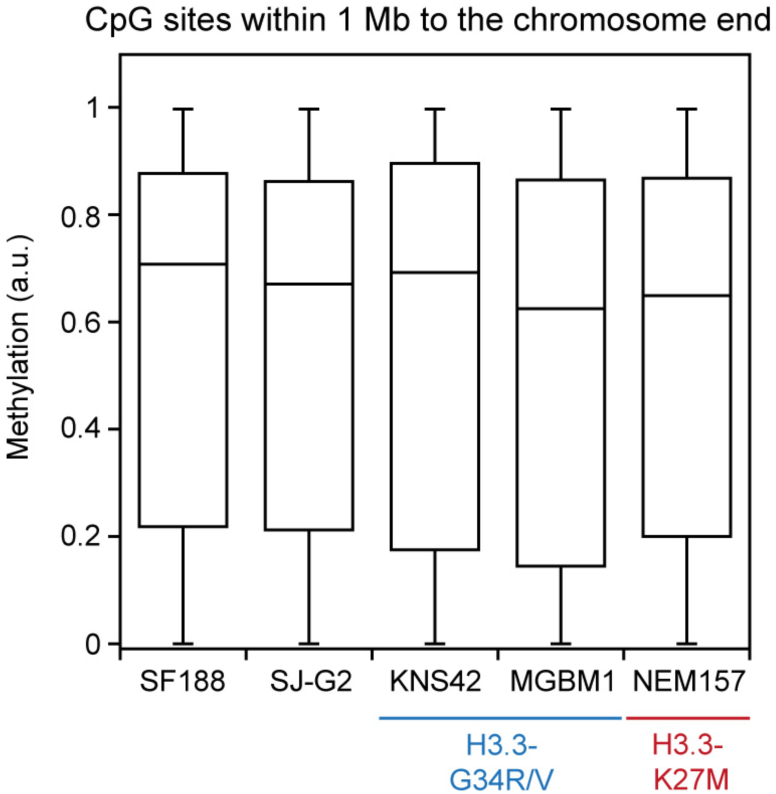
Methylation at CpG sites within one megabase (Mb) to the chromosome end. DNA methylation was determined using the Illumina 450K methylation array. 14,148 CpG sites that were located within 1 Mb from the chromosome end were selected. The median of the corresponding methylation values along with the first and third quartile is shown for each cell line.

**S5 Fig.**
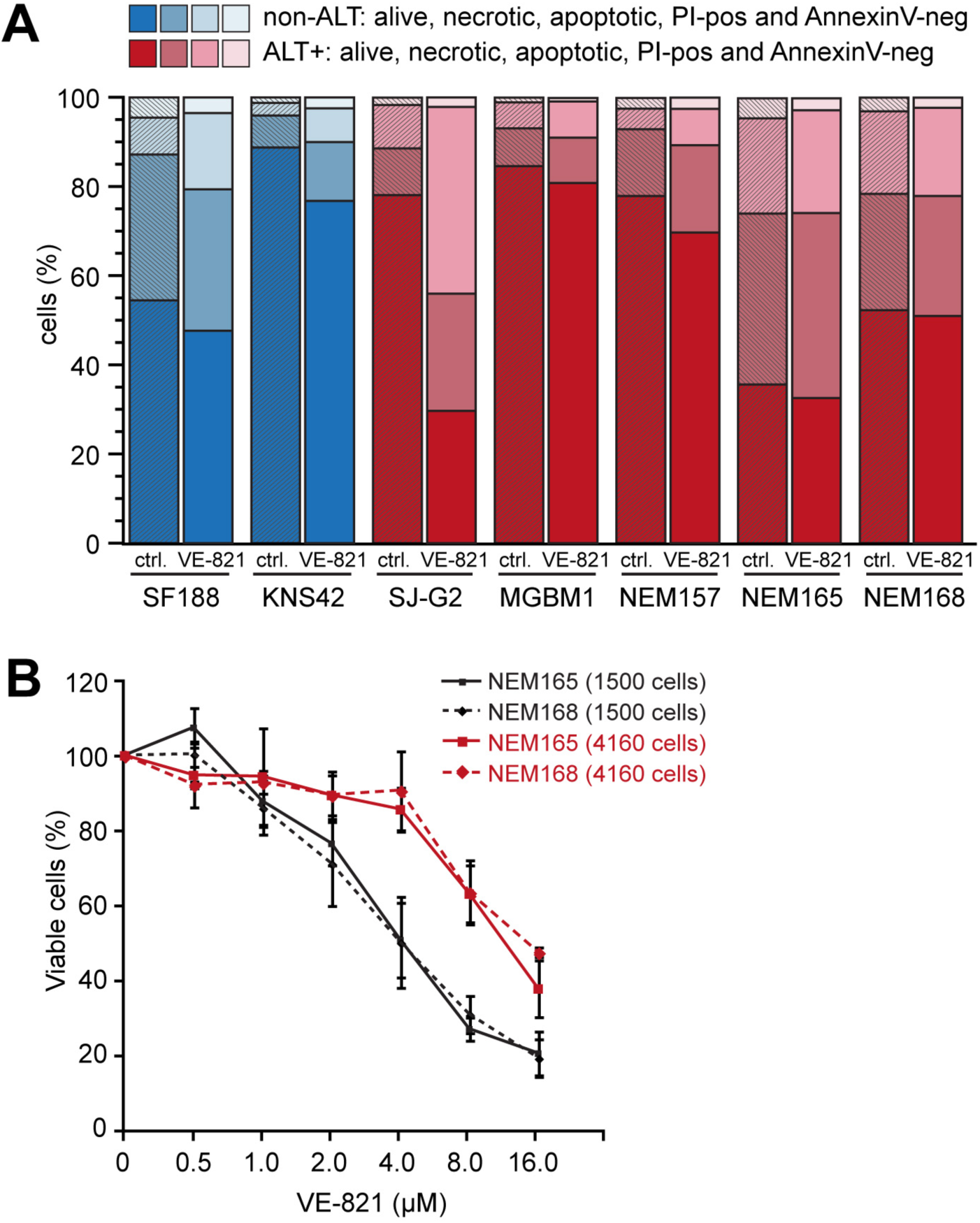
ATRi treatment of pedGBM cell lines. (A) FACS analysis of viable and dead cells in ALT (red color) and non-ALT (blue) cell lines treated with 3 μM VE-821 for 6 days in comparison to the same volume of DMSO. Different cell states were determined by annexin V and propidium iodide staining. The average error was 3.7% ± 0.7% (*n* = 3). (B) A higher starting cell density leads to a lower sensitivity towards the ATR inhibitor VE-821. Different cell numbers of NEM165 and NEM168 were seeded in a 96-well plate and treated with increasing concentrations of the ATR inhibitor VE-821 for 6 days. The starting cell number of 4,160 cells in the 96-well plate corresponds to the cell number used in the FACS experiments. Cell viability was analyzed using the Celltiter Glo assay and is shown as percentage of control (*n* = 3).

**S6 Fig.**
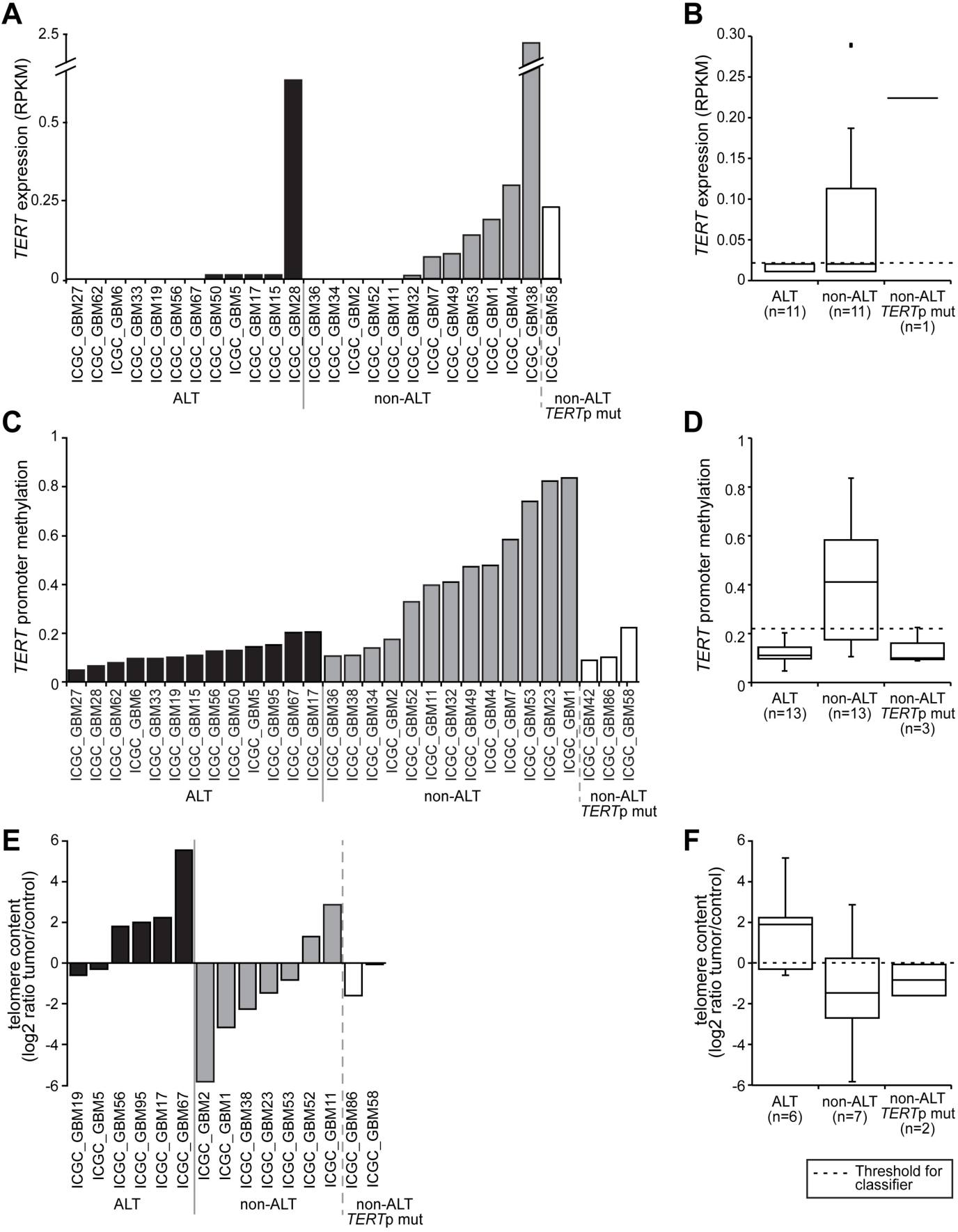
*TERT* mRNA expression, *TERT* promoter methylation, and telomere content in pedGBM samples. Samples were classified as ‘ALT’ (positive result in C-circle assay) and ‘non-ALT’ (either negative results in both C-circle and telomere FISH assays or detection of an activating *TERT* promoter C228T mutation (‘*TERT*p mut’). (A) *TERT* expression extracted from RNA-seq data. (B) Box plot of *TERT* expression data with the threshold for the classifier indicated by a dashed line. Note that outliers >0.5 are not shown. (C) DNA methylation levels of a CpG island upstream of *TERT* that was described to correlate with *TERT* expression [4] as measured by a 450K methylation array. (D) Box plot of DNA methylation levels with the threshold for the classifier indicated by a dashed line. (E) Telomere content as determined by telomere-specific qPCR. The log2 ratio of telomere content measured in the tumor sample to the one in the matched blood control sample is shown. (F) Box plot of telomere content data with the threshold for the classifier indicated by a dashed line.

**S7 Fig.**
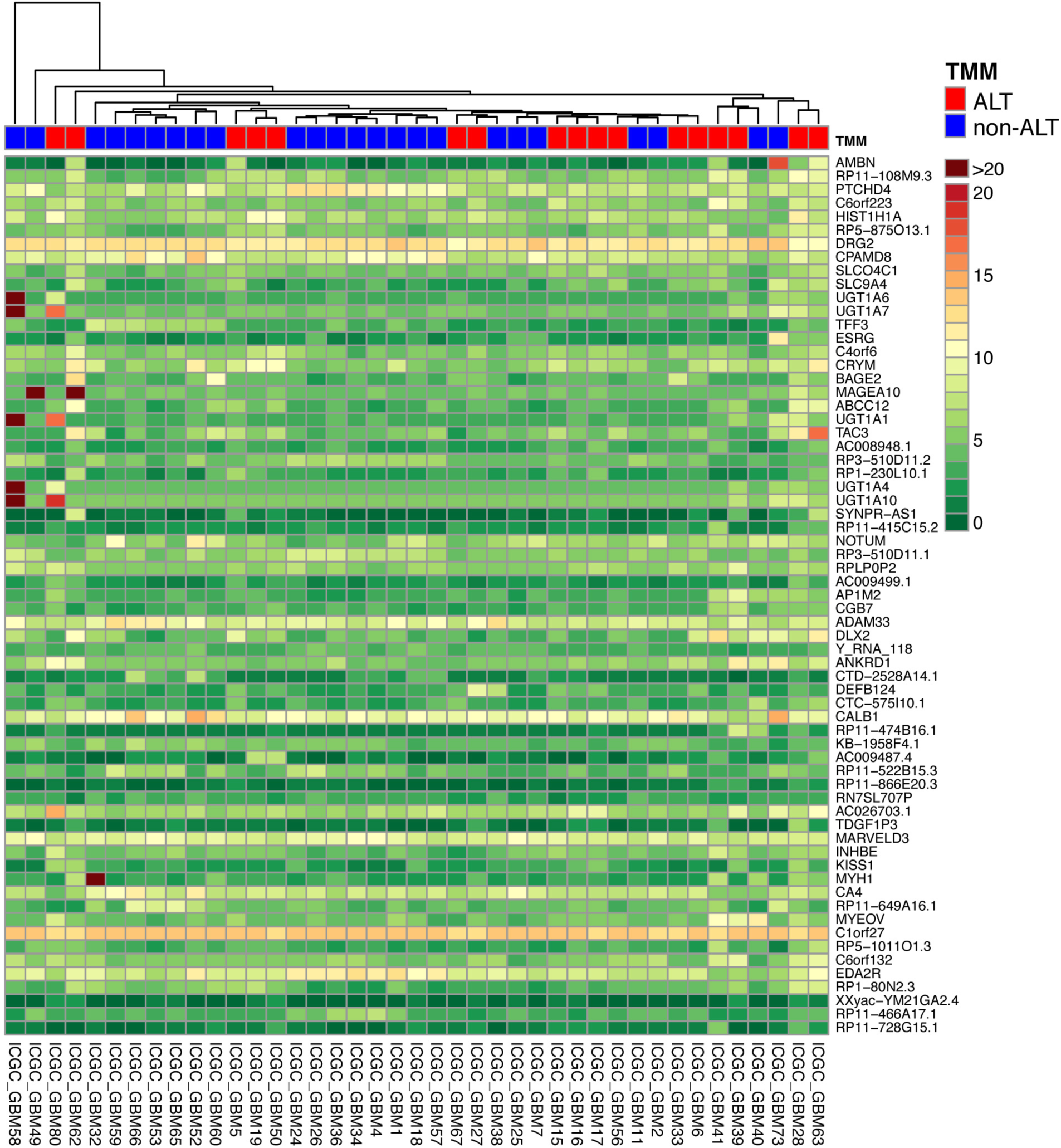
Gene expression analysis of ALT and non-ALT pedGBM samples. A heatmap of the genes that are differentially expressed between ‘ALT’ and ‘non-ALT’ pedGBM samples is shown (adjusted *p*-value ≤ 0.01). The genes were ranked based on significance. The differential expression analysis as well as the normalization was performed with DESeq2. Samples were classified as ‘ALT’ or ‘non-ALT’ using the classification scheme described in the text.

## References

1. Heaphy CM, de Wilde RF, Jiao Y, Klein AP, Edil BH, Shi C, et al. Altered telomeres in tumors with ATRX and DAXX mutations. Science. 2011;333(6041):425. Epub 2011/07/02. doi:10.1126/science.1207313. PubMed PMID: 21719641; PubMed Central PMCID: PMC3174141.

2. Henson JD, Reddel RR. Assaying and investigating Alternative Lengthening of Telomeres activity in human cells and cancers. FEBS Lett. 2010;584(17):3800–11. Epub 2010/06/15. doi:10.1016/j.febslet.2010.06.009. PubMed PMID: 20542034.

3. Dilley RL, Greenberg RA. ALTernative Telomere Maintenance and Cancer. Trends Cancer. 2015;1(2):145–56. doi:10.1016/j.trecan.2015.07.007. PubMed PMID: 26645051; PubMed Central PMCID: PMCPMC4669901.

4. Hakin-Smith V, Jellinek DA, Levy D, Carroll T, Teo M, Timperley WR, et al. Alternative lengthening of telomeres and survival in patients with glioblastoma multiforme. Lancet. 2003;361(9360):836–8. PubMed PMID: 12642053.

5. Schwartzentruber J, Korshunov A, Liu XY, Jones DT, Pfaff E, Jacob K, et al. Driver mutations in histone H3.3 and chromatin remodelling genes in paediatric glioblastoma. Nature. 2012;482(7384):226–31. doi:10.1038/nature10833. PubMed PMID: 22286061.

6. Heaphy CM, Subhawong AP, Hong SM, Goggins MG, Montgomery EA, Gabrielson E, et al. Prevalence of the alternative lengthening of telomeres telomere maintenance mechanism in human cancer subtypes. The American journal of pathology. 2011;179(4):1608–15. Epub 2011/09/06. doi:10.1016/j.ajpath.2011.06.018. PubMed PMID: 21888887; PubMed Central PMCID: PMC3181356.

7. Jiao Y, Killela PJ, Reitman ZJ, Rasheed AB, Heaphy CM, de Wilde RF, et al. Frequent ATRX, CIC, FUBP1 and IDH1 mutations refine the classification of malignant gliomas. Oncotarget. 2012;3(7):709–22. doi:10.18632/oncotarget.588. PubMed PMID: 22869205; PubMed Central PMCID: PMCPMC3443254.

8. Liu X-Y, Gerges N, Korshunov A, Sabha N, Khuong-Quang D-A, Fontebasso AM, et al. Frequent ATRX mutations and loss of expression in adult diffuse astrocytic tumors carrying IDH1/IDH2 and TP53 mutations. Acta neuropathologica. 2012. doi:10.1007/s00401-012-1031-3. PubMed PMID: 22886134.

9. Voon HP, Hughes JR, Rode C, De La Rosa-Velazquez IA, Jenuwein T, Feil R, et al. ATRX plays a key role in maintaining silencing at interstitial heterochromatic loci and imprinted genes. Cell Rep. 2015;11(3):405–18. doi:10.1016/j.celrep.2015.03.036. PubMed PMID: 25865896; PubMed Central PMCID: PMC4410944.

10. Watson LA, Goldberg H, Berube NG. Emerging roles of ATRX in cancer. Epigenomics. 2015;7(8):1365–78. doi:10.2217/epi.15.82. PubMed PMID: 26646632.

11. Rippe K, Luke B. TERRA and the state of the telomere. Nat Struct Mol Biol. 2015;22(11):853–8. doi:10.1038/nsmb.3078. PubMed PMID: 26581519.

12. Elkak A, Mokbel R, Wilson C, Jiang WG, Newbold RF, Mokbel K. hTERT mRNA expression is associated with a poor clinical outcome in human breast cancer. Anticancer Res. 2006;26(6C):4901–4. PubMed PMID: 17214359.

13. Poremba C, Heine B, Diallo R, Heinecke A, Wai D, Schaefer KL, et al. Telomerase as a prognostic marker in breast cancer: high-throughput tissue microarray analysis of hTERT and hTR. J Pathol. 2002;198(2):181–9. doi:10.1002/path.1191. PubMed PMID: 12237877.

14. Lundberg G, Sehic D, Lansberg JK, Ora I, Frigyesi A, Castel V, et al. Alternative lengthening of telomeres--an enhanced chromosomal instability in aggressive non-MYCN amplified and telomere elongated neuroblastomas. Genes Chromosomes Cancer. 2011;50(4):250–62. doi:10.1002/gcc.20850. PubMed PMID: 21319260.

15. Flynn RL, Cox KE, Jeitany M, Wakimoto H, Bryll AR, Ganem NJ, et al. Alternative lengthening of telomeres renders cancer cells hypersensitive to ATR inhibitors. Science. 2015;347(6219):273–7. Epub 2015/01/17. doi:10.1126/science.1257216. PubMed PMID: 25593184.

16. Roth A, Harley CB, Baerlocher GM. Imetelstat (GRN163L)--telomerase-based cancer therapy. Recent Results Cancer Res. 2010;184(0080-0015 (Print)):221–34. doi:10.1007/978-3-642-01222-8_16. PubMed PMID: 20072842.

17. Williams SC. No end in sight for telomerase-targeted cancer drugs. Nat Med. 2013;19(1):6. doi:10.1038/nm0113-6. PubMed PMID: 23295993.

18. Chiappori AA, Kolevska T, Spigel DR, Hager S, Rarick M, Gadgeel S, et al. A randomized phase II study of the telomerase inhibitor imetelstat as maintenance therapy for advanced non-small-cell lung cancer. Ann Oncol. 2015;26(2):354–62. doi:10.1093/annonc/mdu550. PubMed PMID: 25467017; PubMed Central PMCID: PMCPMC4304381.

19. Villa R, Folini M, Lualdi S, Veronese S, Daidone MG, Zaffaroni N. Inhibition of telomerase activity by a cell-penetrating peptide nucleic acid construct in human melanoma cells. FEBS Lett. 2000;473(2):241–8. PubMed PMID: 10812083.

20. Kelland LR. Overcoming the immortality of tumour cells by telomere and telomerase based cancer therapeutics--current status and future prospects. Eur J Cancer. 2005;41(7):971–9. Epub 2005/05/03. doi:10.1016/j.ejca.2004.11.024. PubMed PMID: 15862745.

21. Hu J, Hwang SS, Liesa M, Gan B, Sahin E, Jaskelioff M, et al. Antitelomerase therapy provokes ALT and mitochondrial adaptive mechanisms in cancer. Cell. 2012;148(4):651–63. Epub 2012/02/22. doi:10.1016/j.cell.2011.12.028. PubMed PMID: 22341440; PubMed Central PMCID: PMCPMC3286017.

22. Cesare AJ, Reddel RR. Alternative lengthening of telomeres: models, mechanisms and implications. Nat Rev Genet. 2010;11(5):319–30. Epub 2010/03/31. doi:10.1038/nrg2763. PubMed PMID: 20351727.

23. Nabetani A, Ishikawa F. Alternative lengthening of telomeres pathway: recombination-mediated telomere maintenance mechanism in human cells. J Biochem. 2011;149(1):5–14. Epub 2010/10/13. doi:10.1093/jb/mvq119. PubMed PMID: 20937668.

24. Tokutake Y, Matsumoto T, Watanabe T, Maeda S, Tahara H, Sakamoto S, et al. Extra-chromosomal telomere repeat DNA in telomerase-negative immortalized cell lines. Biochem Biophys Res Commun. 1998;247(3):765–72. Epub 1998/07/02. doi:10.1006/bbrc.1998.8876. PubMed PMID: 9647768.

25. Cesare AJ, Griffith JD. Telomeric DNA in ALT cells is characterized by free telomeric circles and heterogeneous t-loops. Mol Cell Biol. 2004;24(22):9948–57. Epub 2004/10/29. doi:10.1128/MCB.24.22.9948-9957.2004. PubMed PMID: 15509797; PubMed Central PMCID: PMCPMC525488.

26. Henson JD, Cao Y, Huschtscha LI, Chang AC, Au AY, Pickett HA, et al. DNA C-circles are specific and quantifiable markers of alternative-lengthening-of-telomeres activity. Nat Biotechnol. 2009;27(12):1181–5. Epub 2009/11/26. doi:10.1038/nbt.1587. PubMed PMID: 19935656.

27. Chung I, Leonhardt H, Rippe K. De novo assembly of a PML nuclear subcompartment occurs through multiple pathways and induces telomere elongation. J Cell Sci. 2011;124(Pt 21):3603–18. Epub 2011/11/03. doi:10.1242/jcs.084681. PubMed PMID: 22045732.

28. Chung I, Osterwald S, Deeg KI, Rippe K. PML body meets telomere: the beginning of an ALTernate ending? Nucleus. 2012;3(3):263–75. Epub 2012/05/11. doi:10.4161/nucl.20326. PubMed PMID: 22572954; PubMed Central PMCID: PMC3414403.

29. Ng LJ, Cropley JE, Pickett HA, Reddel RR, Suter CM. Telomerase activity is associated with an increase in DNA methylation at the proximal subtelomere and a reduction in telomeric transcription. Nucleic Acids Res. 2009;37(4):1152–9. Epub 2009/01/09. doi:10.1093/nar/gkn1030. PubMed PMID: 19129228; PubMed Central PMCID: PMCPMC2651807.

30. Lovejoy CA, Li W, Reisenweber S, Thongthip S, Bruno J, de Lange T, et al. Loss of ATRX, genome instability, and an altered DNA damage response are hallmarks of the alternative lengthening of telomeres pathway. PLoS Genet. 2012;8(7):e1002772. Epub 2012/07/26. doi:10.1371/journal.pgen.1002772. PubMed PMID: 22829774; PubMed Central PMCID: PMC3400581.

31. Chang FT, Chan FL, JD RM, Udugama M, Mayne L, Collas P, et al. CHK1-driven histone H3.3 serine 31 phosphorylation is important for chromatin maintenance and cell survival in human ALT cancer cells. Nucleic Acids Res. 2015;43(5):2603–14. doi:10.1093/nar/gkv104. PubMed PMID: 25690891; PubMed Central PMCID: PMC4357709.

32. Silvestre DC, Pineda JR, Hoffschir F, Studler J-M, Mouthon M-A, Pflumio F, et al. Alternative lengthening of telomeres in human glioma stem cells. Stem cells (Dayton, Ohio). 2011;29(3):440–51. doi:10.1002/stem.600. PubMed PMID: 21425407.

33. Heaphy CM, Schreck KC, Raabe E, Mao XG, An P, Chu Q, et al. A glioblastoma neurosphere line with alternative lengthening of telomeres. Acta Neuropathol. 2013;126(4):607–8. doi:10.1007/s00401-013-1174-x. PubMed PMID: 24022427; PubMed Central PMCID: PMCPMC3874856.

34. Ran FA, Hsu PD, Wright J, Agarwala V, Scott DA, Zhang F. Genome engineering using the CRISPR-Cas9 system. Nat Protoc. 2013;8(11):2281–308. doi:10.1038/nprot.2013.143. PubMed PMID: 24157548; PubMed Central PMCID: PMCPMC3969860.

35. Arnoult N, Van Beneden A, Decottignies A. Telomere length regulates TERRA levels through increased trimethylation of telomeric H3K9 and HP1alpha. Nat Struct Mol Biol. 2012;19(9):948–56. Epub 2012/08/28. doi:10.1038/nsmb.2364. PubMed PMID: 22922742.

36. O’Callaghan N, Dhillon V, Thomas P, Fenech M. A quantitative real-time PCR method for absolute telomere length. Biotechniques. 2008;44(6):807–9. Epub 2008/05/15. doi:10.2144/000112761. PubMed PMID: 18476834.

37. Cawthon RM. Telomere measurement by quantitative PCR. Nucleic Acids Res. 2002;30(10):e47. PubMed PMID: 12000852; PubMed Central PMCID: PMC115301.

38. Osterwald S, Deeg KI, Chung I, Parisotto D, Wörz S, Rohr K, et al. PML induces compaction, TRF2 depletion and DNA damage signaling at telomeres and promotes their alternative lengthening. J Cell Sci. 2015;128(10):1887–900. doi:10.1242/jcs.148296. PubMed PMID: 25908860.

39. Sturm D, Witt H, Hovestadt V, Khuong-Quang DA, Jones DT, Konermann C, et al. Hotspot mutations in H3F3A and IDH1 define distinct epigenetic and biological subgroups of glioblastoma. Cancer Cell. 2012;22(4):425–37. doi:10.1016/j.ccr.2012.08.024. PubMed PMID: 23079654.

40. Osterwald S, Wörz S, Reymann J, Sieckmann F, Rohr K, Erfle H, et al. A three-dimensional colocalization RNA interference screening platform to elucidate the alternative lengthening of telomeres pathway. Biotechnol J. 2012;7(1):103–16. Epub 2011/06/08. doi:10.1002/biot.201000474. PubMed PMID: 21648092.

41. Wörz S, Sander P, Pfannmöller M, Rieker RJ, Joos S, Mechtersheimer G, et al. 3D Geometry-based quantification of colocalizations in multi-channel 3D microscopy images of human soft tissue tumors. IEEE Trans on Medical Imaging. 2010;29:1474–84. PubMed PMID: 20562043.

42. Deeg KI, Chung I, Bauer C, Rippe K. Cancer Cells with Alternative Lengthening of Telomeres Do Not Display a General Hypersensitivity to ATR Inhibition. Front Oncol. 2016;6:186. doi:10.3389/fonc.2016.00186. PubMed PMID: 27602331; PubMed Central PMCID: PMCPMC4993795.

43. International Cancer Genome Consortium PedBrain Tumor P. Recurrent MET fusion genes represent a drug target in pediatric glioblastoma. Nat Med. 2016;22(11):1314–20. doi:10.1038/nm.4204. PubMed PMID: 27748748.

44. Benjamini Y, Hochberg Y. Controlling the False Discovery Rate - a Practical and Powerful Approach to Multiple Testing. Journal of the Royal Statistical Society Series B-Methodological. 1995;57(1):289–300. PubMed PMID: WOS:A1995QE45300017.

45. Bender S, Tang Y, Lindroth AM, Hovestadt V, Jones DT, Kool M, et al. Reduced H3K27me3 and DNA hypomethylation are major drivers of gene expression in K27M mutant pediatric high-grade gliomas. Cancer Cell. 2013;24(5):660–72. doi:10.1016/j.ccr.2013.10.006. PubMed PMID: 24183680.

46. Grasso CS, Tang Y, Truffaux N, Berlow NE, Liu L, Debily MA, et al. Functionally defined therapeutic targets in diffuse intrinsic pontine glioma. Nat Med. 2015;21(6):555–9. doi:10.1038/nm.3855. PubMed PMID: 25939062; PubMed Central PMCID: PMCPMC4862411.

47. Bjerke L, Mackay A, Nandhabalan M, Burford A, Jury A, Popov S, et al. Histone H3.3. mutations drive pediatric glioblastoma through upregulation of MYCN. Cancer Discov. 2013;3(5):512–9. doi:10.1158/2159-8290.CD-12-0426. PubMed PMID: 23539269; PubMed Central PMCID: PMCPMC3763966.

48. Xue Y, Gibbons R, Yan Z, Yang D, McDowell TL, Sechi S, et al. The ATRX syndrome protein forms a chromatin-remodeling complex with Daxx and localizes in promyelocytic leukemia nuclear bodies. Proc Natl Acad Sci U S A. 2003;100(19):10635–40. PubMed PMID: 12953102.

49. Tang J, Wu S, Liu H, Stratt R, Barak OG, Shiekhattar R, et al. A novel transcription regulatory complex containing death domain-associated protein and the ATR-X syndrome protein. J Biol Chem. 2004;279(19):20369–77. doi:10.1074/jbc.M401321200. PubMed PMID: 14990586.

50. Clynes D, Jelinska C, Xella B, Ayyub H, Scott C, Mitson M, et al. Suppression of the alternative lengthening of telomere pathway by the chromatin remodelling factor ATRX. Nat Commun. 2015;6:7538. doi:10.1038/ncomms8538. PubMed PMID: 26143912.

51. Napier CE, Huschtscha LI, Harvey A, Bower K, Noble JR, Hendrickson EA, et al. ATRX represses alternative lengthening of telomeres. Oncotarget. 2015;6(18):16543–58. doi:10.18632/oncotarget.3846. PubMed PMID: 26001292; PubMed Central PMCID: PMCPMC4599288.

52. Barthel FP, Wei W, Tang M, Martinez-Ledesma E, Hu X, Amin SB, et al. Systematic analysis of telomere length and somatic alterations in 31 cancer types. Nat Genet. 2017:published online 30 January 2017. doi:10.1038/ng.3781. PubMed PMID: 28135248.

53. O’Sullivan RJ, Arnoult N, Lackner DH, Oganesian L, Haggblom C, Corpet A, et al. Rapid induction of alternative lengthening of telomeres by depletion of the histone chaperone ASF1. Nat Struct Mol Biol. 2014;21(2):167–74. doi:10.1038/nsmb.2754. PubMed PMID: 24413054; PubMed Central PMCID: PMC3946341.

54. Nergadze SG, Farnung BO, Wischnewski H, Khoriauli L, Vitelli V, Chawla R, et al. CpG-island promoters drive transcription of human telomeres. RNA. 2009;15(12):2186–94. Epub 2009/10/24. doi:10.1261/rna.1748309. PubMed PMID: 19850908; PubMed Central PMCID: PMC2779677.

55. Hake SB, Garcia BA, Kauer M, Baker SP, Shabanowitz J, Hunt DF, et al. Serine 31 phosphorylation of histone variant H3.3 is specific to regions bordering centromeres in metaphase chromosomes. Proc Natl Acad Sci U S A. 2005;102(18):6344–9. doi:10.1073/pnas.0502413102. PubMed PMID: 15851689; PubMed Central PMCID: PMCPMC1088391.

56. Fujisawa H, Nakajima NI, Sunada S, Lee Y, Hirakawa H, Yajima H, et al. VE-821, an ATR inhibitor, causes radiosensitization in human tumor cells irradiated with high LET radiation. Radiat Oncol. 2015;10:175. doi:10.1186/s13014-015-0464-y. PubMed PMID: 26286029; PubMed Central PMCID: PMCPMC4554350.

57. Sanjiv K, Hagenkort A, Calderon-Montano JM, Koolmeister T, Reaper PM, Mortusewicz O, et al. Cancer-Specific Synthetic Lethality between ATR and CHK1 Kinase Activities. Cell Rep. 2016;14(2):298–309. doi:10.1016/j.celrep.2015.12.032. PubMed PMID: 26748709; PubMed Central PMCID: PMCPMC4713868.

58. Heidenreich B, Rachakonda PS, Hemminki K, Kumar R. TERT promoter mutations in cancer development. Curr Opin Genet Dev. 2014;24:30–7. doi:10.1016/j.gde.2013.11.005. PubMed PMID: 24657534.

59. Stern JL, Theodorescu D, Vogelstein B, Papadopoulos N, Cech TR. Mutation of the TERT promoter, switch to active chromatin, and monoallelic TERT expression in multiple cancers. Genes Dev. 2015;29(21):2219–24. doi:10.1101/gad.269498.115. PubMed PMID: 26515115; PubMed Central PMCID: PMCPMC4647555.

60. Killela PJ, Reitman ZJ, Jiao Y, Bettegowda C, Agrawal N, Diaz LA, Jr., et al. TERT promoter mutations occur frequently in gliomas and a subset of tumors derived from cells with low rates of self-renewal. Proc Natl Acad Sci U S A. 2013;110(15):6021–6. doi:10.1073/pnas.1303607110. PubMed PMID: 23530248; PubMed Central PMCID: PMCPMC3625331.

61. Mangerel J, Price A, Castelo-Branco P, Brzezinski J, Buczkowicz P, Rakopoulos P, et al. Alternative lengthening of telomeres is enriched in, and impacts survival of TP53 mutant pediatric malignant brain tumors. Acta Neuropathologica. 2014;128(6):853–62. doi:10.1007/s00401-014-1348-1. PubMed PMID: 25315281.

62. Bryan TM, Englezou A, Dalla-Pozza L, Dunham MA, Reddel RR. Evidence for an alternative mechanism for maintaining telomere length in human tumors and tumor-derived cell lines. Nat Med. 1997;3(11):1271–4. PubMed PMID: 9359704.

63. Henson JD, Neumann AA, Yeager TR, Reddel RR. Alternative lengthening of telomeres in mammalian cells. Oncogene. 2002;21(4):598–610. doi:10.1038/sj.onc.1205058. PubMed PMID: 11850785.

64. Ernst A, Jones DT, Maass KK, Rode A, Deeg KI, Jebaraj BM, et al. Telomere dysfunction and chromothripsis. Int J Cancer. 2016;138(12):2905–14. doi:10.1002/ijc.30033. PubMed PMID: 26856307.

65. Maciejowski J, Li Y, Bosco N, Campbell PJ, de Lange T. Chromothripsis and Kataegis Induced by Telomere Crisis. Cell. 2015;163(7):1641–54. doi:10.1016/j.cell.2015.11.054. PubMed PMID: 26687355; PubMed Central PMCID: PMCPMC4687025.

66. Ducrest AL, Amacker M, Mathieu YD, Cuthbert AP, Trott DA, Newbold RF, et al. Regulation of human telomerase activity: repression by normal chromosome 3 abolishes nuclear telomerase reverse transcriptase transcripts but does not affect c-Myc activity. Cancer Res. 2001;61(20):7594–602. PubMed PMID: 11606399.

67. Fredriksson NJ, Ny L, Nilsson JA, Larsson E. Systematic analysis of noncoding somatic mutations and gene expression alterations across 14 tumor types. Nat Genet. 2014;46(12):1258–63. doi:10.1038/ng.3141. PubMed PMID: 25383969.

68. Castelo-Branco P, Choufani S, Mack S, Gallagher D, Zhang C, Lipman T, et al. Methylation of the TERT promoter and risk stratification of childhood brain tumours: an integrative genomic and molecular study. Lancet Oncol. 2013;14(6):534–42. doi:10.1016/S1470-2045(13)70110-4. PubMed PMID: 23598174.

69. Sturm D, Bender S, Jones DT, Lichter P, Grill J, Becher O, et al. Paediatric and adult glioblastoma: multiform (epi)genomic culprits emerge. Nat Rev Cancer. 2014;14(2):92– 107. doi:10.1038/nrc3655. PubMed PMID: 24457416; PubMed Central PMCID: PMCPMC4003223.

70. Heaphy CM, Subhawong AP, Hong SM, Goggins MG, Montgomery EA, Gabrielson E, et al. Prevalence of the alternative lengthening of telomeres telomere maintenance mechanism in human cancer subtypes. Am J Pathol. 2011;179(4):1608–15. Epub 2011/09/06. doi:10.1016/j.ajpath.2011.06.018. PubMed PMID: 21888887; PubMed Central PMCID: PMCPMC3181356.

71. Taylor KR, Mackay A, Truffaux N, Butterfield YS, Morozova O, Philippe C, et al. Recurrent activating ACVR1 mutations in diffuse intrinsic pontine glioma. Nat Genet. 2014;46(5):457–61. doi:10.1038/ng.2925. PubMed PMID: 24705252; PubMed Central PMCID: PMCPMC4018681.

72. Wu G, Diaz AK, Paugh BS, Rankin SL, Ju B, Li Y, et al. The genomic landscape of diffuse intrinsic pontine glioma and pediatric non-brainstem high-grade glioma. Nat Genet. 2014;46(5):444–50. doi:10.1038/ng.2938. PubMed PMID: 24705251; PubMed Central PMCID: PMCPMC4056452.

73. Leung JW, Ghosal G, Wang W, Shen X, Wang J, Li L, et al. Alpha thalassemia/mental retardation syndrome X-linked gene product ATRX is required for proper replication restart and cellular resistance to replication stress. J Biol Chem. 2013;288(9):6342–50. doi:10.1074/jbc.M112.411603. PubMed PMID: 23329831; PubMed Central PMCID: PMC3585069.

74. McDonald KL, McDonnell J, Muntoni A, Henson JD, Hegi ME, von Deimling A, et al. Presence of alternative lengthening of telomeres mechanism in patients with glioblastoma identifies a less aggressive tumor type with longer survival. J Neuropathol Exp Neurol. 2010;69(7):729–36. Epub 2010/06/11. doi:10.1097/NEN.0b013e3181e576cf. PubMed PMID: 20535033.

75. Gunkel M, Chung I, Worz S, Deeg KI, Simon R, Sauter G, et al. Quantification of telomere features in tumor tissue sections by an automated 3D imaging-based workflow. Methods. 2016:published online 7 October 2016. doi:10.1016/j.ymeth.2016.09.014. PubMed PMID: 27725304.

## Supplementary References

76. Bender S, et al., International Cancer Genome Consortium PedBrain Tumor Project (2016) Recurrent MET fusion genes represent a drug target in pediatric glioblastoma. Nat Med: published online 17 October 2016 Doi 10.1038/nm.4204

77. Bender S, Tang Y, Lindroth AM, Hovestadt V, Jones DT, Kool M, Zapatka M, Northcott PA, Sturm D, Wang W et al (2013) Reduced H3K27me3 and DNA hypomethylation are major drivers of gene expression in K27M mutant pediatric high-grade gliomas. Cancer Cell 24: 660-672 Doi 10.1016/j.ccr.2013.10.006

78. Benjamini Y, Hochberg Y (1995) Controlling the False Discovery Rate - a Practical and Powerful Approach to Multiple Testing. J Roy Stat Soc B Met 57: 289–300

79. Castelo-Branco P, Choufani S, Mack S, Gallagher D, Zhang C, Lipman T, Zhukova N, Walker EJ, Martin D, Merino Det al (2013) Methylation of the TERT promoter and risk stratification of childhood brain tumours: an integrative genomic and molecular study. Lancet Oncol 14: 534–542 Doi 10.1016/S1470-2045(13)70110-4

80. Love MI, Huber W, Anders S (2014) Moderated estimation of fold change and dispersion for RNA-seq data with DESeq2. Genome Biol 15: 550 Doi 10.1186/s13059-014-0550-8

